# RAMZIS: a bioinformatic toolkit for rigorous assessment of the alterations to glycoprotein structure that occur during biological processes

**DOI:** 10.1101/2023.05.30.542895

**Authors:** William Edwin Hackett, Deborah Chang, Luis Carvalho, Joseph Zaia

**Affiliations:** Boston University, Bioinformatics Program, One Silber Way, Boston 02215, MA, USA; Department of Biochemistry, Boston University, One Silber Way, Boston 02215; Department of Mathematics, Boston University, One Silber Way, Boston 02215

## Abstract

**Motivation:** Glycosylation elaborates the structures and functions of glycoproteins; glycoproteins are common post-translationally modified proteins and are heterogeneous and non-deterministically synthesized as an evolutionarily driven mechanism that elaborates the functions of glycosylated gene products. While glycoproteins account for approximately half of all proteins, their macro- and micro-heterogeneity requires specialized proteomics data analysis methods as a given glycosite can be divided into several glycosylated forms, each of which must be quantified. Sampling of heterogeneous glycopeptides is limited by mass spectrometer speed and sensitivity, resulting in missing values. In conjunction with the low sample size inherent to glycoproteomics, this necessitated specialized statistical metrics to identify if observed changes in glycopeptide abundances are biologically significant or due to data quality limitations.

**Results:** We developed an R package, Relative Assessment of *m/z* Identifications by Similarity (RAMZIS), that uses similarity metrics to guide biomedical researchers to a more rigorous interpretation of glycoproteomics data. RAMZIS uses contextual similarity to assess the quality of mass spectral data and generates graphical output that demonstrates the likelihood of finding biologically significant differences in glycosylation abundance dataset. Investigators can assess dataset quality, holistically differentiate glycosites, and identify which glycopeptides are responsible for glycosylation pattern expression change. Herein RAMZIS approach is validated by theoretical cases and by a proof-of-concept application. RAMZIS enables comparison between datasets too stochastic, small, or sparse for interpolation while acknowledging these issues in its assessment. Using our tool, researchers will be able to rigorously define the role of glycosylation and the changes that occur during biological processes.

**Availability:** https://github.com/WillHackett22/RAMZIS

**Contact:** Joseph Zaia, Boston University Medical Campus, 670 Albany St., rm 509, Boston, MA 02118 USA, (e), (v) 1-617-358-2429

**Supplementary information:** Supplementary data are available

## 1. Introduction

### 1.1 Biological Context of Glycoproteins

Evolutionary pressure for multicellularity and pathogen defense drives the complexity of protein glycosylation [1]. Mammalian glycoprotein glycosites are modified with a distribution of glycoforms resulting from the non-template driven ER-Golgi biosynthesis. Glycosylation modulates protein physicochemical properties and the interactions with protein binding partners, and each glycosylation site on a protein contributes to these changes. To understand the biological functions of individual glycoproteins, it is necessary to determine the quantitative changes to protein site glycosylation. This remains a challenge because the glycan heterogeneity at each glycosite multiplies the number of precursor ions that must be sampled relative to a classical proteomics experiment; the ability to sample these heterogeneous glycopeptide glycoforms is limited by analyzer speed. On the one hand, data dependent acquisition (DDA) of glycopeptides results in a missing values problem the severity of which increases with the number of co-eluting glycoforms and decreasing glycoform abundance [2]. On the other, data independent acquisition (DIA) samples all precursor ions but sensitivity is limited by the need to scan windows over the mass range [3]. As a result, the ability to quantify changes in glycoprotein glycosylation is limited by sampling deficiencies.

For a given glycosite, the glycopeptide glycoforms elute over a narrow reversed-phase chromatography retention time range. Quantification of these glycoforms is based on their extracted ion chromatograms over a 2- to 3-minute window, but because many glycoforms may be present at each glycosite, some which will be missed in any given replicate simply due to the acquisition limitations of tandem mass spectrometry, MS/MS. Confident identification and quantification of glycopeptide glycoforms requires tandem MS acquisition of the glycopeptide precursor ions [4-5]. The statistical power of the quantitative results is limited by the number of glycoforms for a given glycosite that are missed in any given sample replicate.

As shown by numerous recent publications and preprints concerning the SARS-CoV-2 spike protein [6-12], there are many biomedical laboratories with the capability to assign site-specific glycosylation for highly complex glycoproteins using DDA LC-MS/MS datasets. For such experiments, the number of glycopeptide glycoforms observed in the MS1 dimension may naively be regarded as complete. However, if the MS1 dimension is used as the sole measure for assigning glycopeptides, the confidence of assignments would be low, owing to the lack of tandem MS confirmation of glycopeptide assignments. The confidence would be higher if the tandem MS acquisition targets the most abundant subset of the glycopeptide glycoforms as in DDA LC-MS acquisition experiments used by the majority of published glycoproteomics experiments in the recent literature [6-8,10-13]. Those glycoforms not subjected to tandem MS can be inferred based on exact mass from the MS1 dimension only with significant false assignments resulting from co-eluting glycopeptides.

The challenge is therefore to assign the appropriate degree of confidence for quantitative results for DDA datasets. In principle, as the number of glycosites in a proteolytic digest increases, the likelihood of overlapping precursor mass and retention time values increases. The user is left needing to judge whether an observed difference in glycopeptide abundances is likely to be biologically significant. To address this need we developed a guiding metric for users in glycoproteomics quantification.

### 1.2 Statistical Context of Glycoproteins

Conventional statistics, such as t-tests and ANOVAs, are unlikely to provide accurate results in glycoproteomics datasets as they rely on a greater number of replicates, a lower degree of missingness, and an assumption of relative independence. This becomes especially true when attempting to compare quantifications for every single observed glycopeptide at every glycosite, as the likelihood of false positives increases in proportion to the number of tests performed in a given dataset. While log-transformed glycopeptide abundances are normally distributed on a population level, that does not mean that individual log-transformed abundances are normally distributed. Most glycoproteomic studies cannot prove normality due to the paucity of sample size; sample sizes greater than 30 can generally be assumed normal, but glycoproteomic experiments often do not have 30 samples in total, let alone for each sample group. While normality can be tested for below this threshold, the likelihood of identifying a real deviation from normal is low due to the power of normality tests. As sample size is a continuous, inherent problem in glycoproteomic experimental design, this, in addition to the interdependence of glycopeptide quantitation, removes standard parametric testing as a viable option. Many parametric tests discard quantification information in favor of ranking, which reduces our ability to account for effect sizes and has lower power than is acceptable for our typical sample sizes. Fortunately, datasets may be compared as collectives, trading specificity for power.

Similarity measures indicate how much two things resemble one another and are used in clustering algorithms and sequence alignment [14-15]. Chemoinformatics, the joint field of computer science applied to physical chemistry, makes use of them in molecular fingerprinting, a tool for drug screening and chemical mapping [16]. The Tanimoto coefficient has been used to evaluate biological species co-occurrences based on presence/absence data to elucidate relationships among organisms [17]. The Tanimoto similarity metric identifies a molecule by comparison to a database based on a set of chemical and physical attributes. A Tanimoto similarity metric transforms the various attributes of the molecule to a simple scale of 0, not at all similar, to 1, practically identical.

Glycoproteomic relative abundances can be grouped by glycosite and compared using this transformation, but additional scaling factors should be introduced to account for the missingness in glycoproteomics data and its adverse influence on reliability. We weighed the Tanimoto similarity metric for this purpose. We also developed contextual similarity, a form of permutation test wherein a sampling of data points provides the context needed to understand the results of similarity comparisons.

Three types of comparison provide the user with a better understanding of the analysis. First, the quality of a dataset and outlier identification is observed through a Self or Internal comparison. Second, the desired comparisons of control versus variant (mutant or disease state), here referred to as Test comparisons, provide a sampling of the desired comparison. And third, a null hypothesis is generated and comparisons made as if there were no difference between the two samples. This is simulated by comparing random samplings of a joint dataset; this shows the similarity behavior wherein glycopeptides from both sample datasets were sampled from the same underlying distribution. Combined with the Test comparisons, the Null hypothesis indicates the likelihood of separable, differentiable glycosites, and informs on the viability of the comparison. After providing a general assessment of viability, we then rank individual glycopeptides for follow up targeted experiments or more conventional statistical analyses. Here, we describe the RAMZIS internal algorithm and criterion and its application to re-analysis of published data, reconfirming prior work, and expanding on these analyses.

## 2. Methods

RAMZIS is an R package that takes the abundances of two sample group glycopeptides as input and produces a data quality assessment and visualization, a general comparison with a visualization, and a ranked list of glycopeptides as output. It does this via contextual similarity, whereby related similarity comparisons made with a modified Tanimoto similarity metric provide context for one another as a form of permutation test. The overall algorithm is illustrated in Supplemental Figure 1.

### 2.1 Modified Tanimoto Metric

The Tanimoto similarity metric was modified in order to increase our ability to differentiate glycoprotein glycosites based on abundances of glycopeptides. The traditional metric is the sum of sample group A times sample group B scaled to the contributions of A and B as follows:

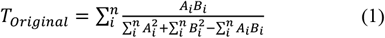

We modified the metric to include both the presence and abundance of glycopeptide glycoforms as follows:

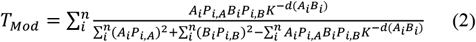

Where i is an identification; n is the number of identifications in a subset or glycosite; A and B refer to sample groups, meaning Ai refers to the relative log abundance of glycopeptide i in group A; Pi,A refers to the presence of glycopeptide i in group A; K is a distance scaling term equal to 1 plus the mean presence rate of glycopeptide i in both groups; d(Ai,Bi) is the Manhattan distance between both groups at glycopeptide i.

These modifications were developed through heuristic observation based on data that compare site-specific glycosylation of glycoproteins with known quality and comparability differences.

### 2.2 RAMZIS Workflow

#### 2.2.1 Step 1: Data Format & Standardization

The input for RAMZIS is two matrices of the relative abundances of identified glycopeptides by sample; it can delineate between exact peptide matches, and glycosite notations can be given to subset the data more comprehensively.

RAMZIS modulates signal strength to be on the same scale through relative log abundance. Optionally, it can also scale in proportion to the log of the provided TIC of each replicate. This ensures that the replicate with the least overall signal does not over-represent an individual glycopeptide due to fewer identifications to compare against. A minimum number of observations per identification is used to pre-emptively filter poor quality results, which can be disabled as desired.

#### 2.2.2 Step 2: Bootstrapping Datasets

As described earlier replicates were sampled with replacement with replacement to produce three kinds of collections of datasets that were then used to produce four similarity distributions. A collection herein refers to the set of all sampled datasets of a specific type.

A test collection is composed of multiple test datasets, which are each made solely from one of the sample groups with one fewer replicate than the observed sample group. This is done for both sample groups, making two test collections. For each test collection, 100 samplings are created prior to duplicate reduction. The test sets are used to generate two Internal Similarity Distributions and the Test Similarity Distribution.

A null collection is made of datasets that sample from both sample groups with replicates equal to the average number of replicates seen in the original sample groups. Sampling is done at a roughly equal rate from both groups to prevent outlier effects; duplicate datasets are eliminated from the collection to prevent spurious outlier effects in the similarity comparisons. For each null set, 200 datasets are generated prior to duplicate reduction. The null collection is split into two null collections with no sampling overlaps to be tested against one another.

#### 2.2.3 Step 3: Similarity Distributions and Observed Similarity

Five groups of similarity comparison are made. One is the test collection comparison, which calculates the similarity between every sampling from one test collection to every sampling of the other test collection; this creates a maximum of ten thousand test comparisons. These test comparisons make up the Test Similarity Distribution; it simulates the extent to which the two sample groups resemble one another.

The second and third groups of similarity comparisons are the internal comparisons, which compare samplings of a sample group’s test collection against the rest of that group’s test collection. This generates a maximum of 4950 internal comparisons, limited to prevent combinatorial expansion in larger datasets. These distributions are the Internal Similarity Distributions for their respective sample groups; they represent the extent to which a sample group resembles itself and provide a check on data quality through internal consistency.

The fourth group of similarity comparisons are null comparisons. These work like the test comparisons. They produce a maximum of 40,000 comparisons, limited to prevent combinatorial expansion and reduce computation needed. This distribution of comparisons is the Null Similarity Distribution; it represents how much a random comparison within the dataset resembles any other random comparison within the dataset. It is used as a reference point for the Test Similarity Distribution.

The final similarity comparison grouping is called the Observed Similarity Comparison and is the singular comparison between the observed sample groups from the original data. This is the similarity of the data without bootstrapping and is a key reference value in evaluating simulation quality, data quality, and glycopeptide ranking.

#### 2.2.4 Step 4: General Data Quality Assessment

The extent to which similarity comparisons can be made meaningfully is influenced by the quality of the LC-MS data ion abundances and reproducibility. The similarity distributions allow for assessing this quality through their means, variances, and modalities.

The first assessment uses the two Internal Distributions. This is done with two intents: to determine if there are outliers in the data and to determine the reliability of the data. Multimodal distributions have their peak membership tested. Peak membership refers to the set of replicates involved in the comparison; non-uniform peak membership indicates an outlier effect.

Deviation from uniform distribution is determined via a z-score, where the replicate in question is withheld from the mean and standard deviation calculations. If the z-score exceeds 3 for a sample in a peak then it is marked as being either over or underrepresented within that peak; it should only be removed if it is under-represented in the primary peak or overrepresented in a non-primary peak, and if that peak contains more than 10% of the overall comparisons. RAMZIS is currently unable to use this outlier detection method for subgroup identification due to the number of subset comparisons required to test this. If the distributions are still multimodal after performing replicate removal, then there is either an underlying data quality problem or a subset within the sample group; either will prevent appropriate comparison in the given dataset.

Internal Distributions are also assessed via a confidence score. This score is the average weighted z-score defined in equation 5; a score of 2 indicates less than a 5% chance of overlap between the Internal and Test Distribution means; more conservative users can raise this threshold as they feel appropriate. The score is the absolute distance between the means of the Internal and Test Similarities, divided by the standard deviation of the Internal Similarity, and then scaled according to the degree of overlap between the two distributions. The overlap scaling term is the sum of the false positive rate, alpha, and the false negative rate, beta. Alpha and Beta overlaps are further explained in the Supplemental Text and generally defined by equations 3 and 4:

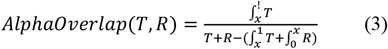

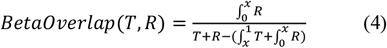

Where T is the density of the Test Distribution; R is the density of the reference similarity distribution, in this case the Internal Similarities; x is the point at which the Test begins to have a lower density than the reference.

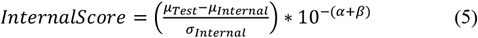

Internal Distributions will have different Internal Confidence (IC) scores for each comparison they are used in. A prior version of the score was used in a related study [8]; that version used the ratio of the internal peak height to the variance, scaled according to an overlap weighting term. This left out information relating to the distance between Similarity distributions; reanalysis did not substantially change the prior analysis.

The mean similarity of the Internal Distributions is only relevant in the context of the Test Similarity. The data can only be considered of sufficient quality if both Internal Distributions have means greater than the mean of the Test Distribution. An Internal Similarity lower than the Test by more than one of either of their standard deviations indicates that a sample group has a higher degree of variability compared to itself, than it does when compared to an outside entity, which indicates the sample group cannot be viewed as a unified group in the context of a comparison to the other sample group. Further analysis should treat the low Internal Similarity group as individual data points rather than replicates. Internal Distributions equal to the Test Distribution indicate a subset, further explained in the Supplemental Text.

The Observed Similarity must be within 3 standard deviations of the Test Distribution’s mean for it to be considered a typical comparison. A higher standard can be imposed by only being considered typical if it is within the central 50% of the Test Distribution. Typicality is used as a stand-in for assessing the bootstrapping; if the Observed Similarity is atypical for the Test Distribution then the Test Distribution cannot give us a reliable assessment of the data.

A Null Distribution lower than its Test Distribution indicates very high degrees of internal variability and low degrees of missingness; this would likely be identified in the Internal Distributions.

#### 2.2.5 Step 5: General Comparison

If data quality is acceptable and outlier effects are removed, a general comparison between the Test and Null Distributions is made. If the two underlying distributions are significantly differentiable then it is likely that the sample groups have differentiable quantification patterns. Differentiability is determined here by overlap proportion as described above in equations 3 and 4. Conventional standards place differentiability at an alpha less than 0.05 and a beta less than 0.20.

If alpha and beta are below their thresholds, then the two sample groups are considered differentiable in their overall quantification patterns. The ability to globally differentiate the groups does not guarantee the existence of a consistent differentiable identification between the groups, but it does make it more likely. It is possible that individual assessments will indicate a need for improved data quality rather than be differentiable themselves. This overview tells the user that differentiating the sample groups is possible.

A visual display is generated to assist the user in understanding this comparison. It displays the density of the Test Distribution in red, the density of the Null Distribution in blue, the false positive region in black, and the false negative region in grey. Different examples are presented in the results section.

#### 2.2.6 Step 6: Ranking Assessment and Reporting

To guide analysis and experimental design, individual glycopeptide identifications are ranked based on their contributions to the various similarity distributions. Contribution refers to the amount of similarity an identification adds to the total similarity in each comparison. Individual identifications undergo a quality assessment by their contribution to Internal Similarity by examining peak membership and relative position to their Test Similarity contribution. These assessments are performed as they are for the totaled distributions, but combine alpha and beta due to resolution for a total threshold of 25%. Potential causes of failure are identified and reported alongside the identifications rankings.

Rank is correlated to the differentiation between their Null and Test group contributions. Unlike the quality assessment, the contributions used herein only use the numerator term of equation 2; this is because missing values bias the overall denominator term and more likely to occur in Null type datasets, ironically preventing the identification of missing values as contributors to differences in similarity. For a given identification, these unweighted contributions use the average z-score of all Test contributions from the context of the Null distribution. Therefore, the lower the z-score, the higher the ranking. Ideal ranked glycopeptides will be less than -2 if they are sources of differentiation, but this number may be higher in cases of missing values. Significant positive z-scores may indicate an underlying interdependence between datasets.

## 3 Results

To display the workings and utility of RAMZIS for a glycoproteomic researcher and explain the usage, we provide examples and a proof-of-concept reexamination. These come in the form of theoretical abundance data as a guide and exploration of published experimental data. The guide provides a range of plausible scenarios to better illustrate the output and utility of the toolset and consists of Supplemental Figures 2-10 and Supplemental Tables 2-4. The reexamination takes data from a published glycoproteomics study [18] and expands its analysis.

### 3.1 AGP Case Study Premise

A prior experiment found that as sample heterogeneity increases, the ability to identify and quantify glycopeptides decreases [18]. This problem can be alleviated using an informed search space, one that uses proteomic and glycomic data to create a more targeted search space. These informed search spaces increase the confidence of our results and the likelihood of identifying and quantifying glycopeptides. The prior experiment looked at purified alpha-1-acid glycoprotein (AGP), as well as AGP spiked into increasing amounts of human serum glycoproteins.

These data were analyzed using a standard GlycReSoft analysis workflow [8] using default settings. Peaks Studio 8 [19] was used for the proteomics search. GlycReSoft was used for the glycan search. Both the glycomics and proteomics databases were used to generate the informed search spaces. The naive search spaces that were used to compare against the informed search space were combinatorial glycan search spaces attached to all proteins expected to be in the mixture. All searches were performed at 5% FDR and used fixed carbamidomethylation and variable oxidation, deamidation, and pyro-glutamination.

#### 3.1.1 AGP: Naive vs Informed in a Homogeneous Mixture

In glycoproteomics studies, it is tempting to make assumptions about the glycoprotein purity. We showed that it is better to build a glycoproteomics search space based on measured glycome and proteome information [20]. A naïve search space uses sets of glycan compositions from public databases and assumes that the target glycoprotein is pure. An informed search space uses measured glycomes and proteomes to define the set of theoretical glycopeptides used to search the data.

We applied RAMZIS to the published data from that study and found that, even in a low complexity mixture of purified AGP, we identified differences in the overall glycosylation pattern between naïve and informed search spaces as shown in Figure 1. This analysis focuses on the 4th AGP glycosite, herein YNTT. RAMZIS identified an outlier sample from this dataset, reducing the sample size to three separate LC runs; this outlier identification and removal process can be found in Supplemental Figure 11. The remaining multimodality is due to a higher quality replicate type outlier.

**Figure 1.**
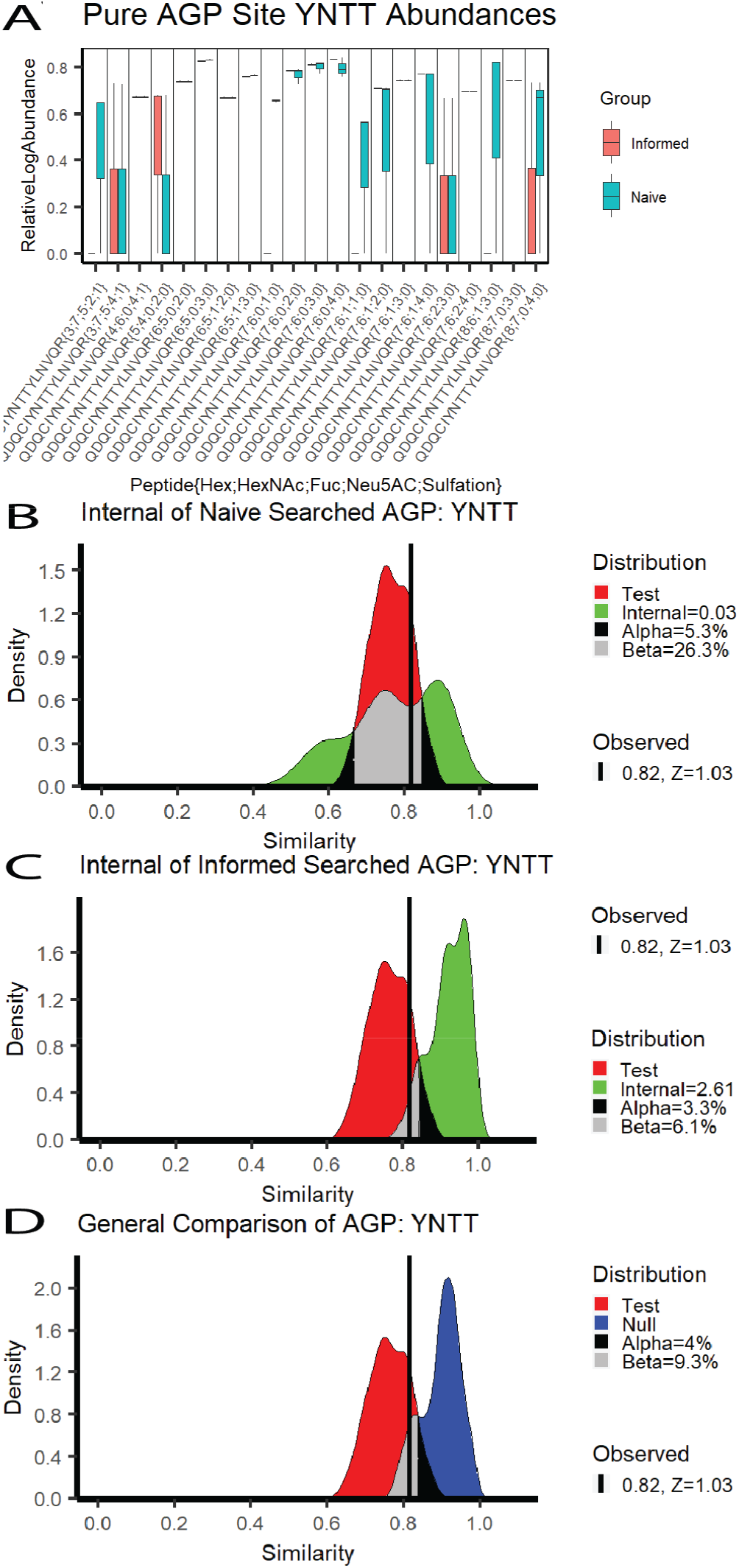
AGP Naïve vs. Informed Search Space Comparison. A) The relative log abundances of the glycopeptides observed in pure AGP samples; the Informed search values are in red, and the Naïve search values are in teal; these have been standardized by TIC. B) The Internal Similarity of the Naive search space with IC=0.03, FPR=5.3%, FNR=26.3%. C) The Internal Similarity of the Naive search results with IC=2.61, FPR=3.3%, FNR=6.1%. D) The general comparison between the Test and Null similarities of the Naive and Informed search space results with FNR=4%, FPR=9.3%, and has the observed similarity at 0.82 with a Test Similarity z-score of 1.03

In Figure 1a, the standardized log abundances of the glycopeptides are displayed in a pairwise boxplot, with the Informed search results in red on the left and the Naïve in teal on the right. The quantifications appear highly consistent with notable visual exceptions. Boxplots such as the abundances at YNNT{3;7;5;4;1} indicate that two of three samples are zero, with the third at the line end point.

Figures 1b and 1c are both Internal similarity plots with the Test similarity in red and the Internal similarities in green. Figure 1b shows the Naïve search’s Internal Similarity with an IC of 0.03, an FPR of 5.3%, and an FNR of 26.3%. These fail all quality metrics, which indicates that the Naïve has insufficient quality for a comparison against the Informed. Figure 1c shows the Internal similarity of the Informed search with an IC of 2.61, an FPR of 3.3%, and an FNR of 6.1%. These pass all quality metrics, indicating that the Informed has a high enough data quality for this comparison. The Test similarity meets its own quality threshold with the observed similarity of the original comparison at 0.82 with a z-score of 1.03, less than 3.

A truncated, reformatted ranking data summary from RAMZIS. Each row shows a different glycopeptide’s identity (PEP{Hex;HexNAc;Fuc;Neu5Ac;Sulfate}), contribution z-score, quality indicator, and probable quality failure cause.

The general similarity comparison is shown in Figure 1d, comparing the Test to the Null similarity distributions. The Null is in blue, and there is a 4% FPR and a 9.3% FNR. Without the data quality problems seen in Naïve, the glycosylation patterns could be considered differentiable, but. The glycopeptides that are ranked to be the most likely contributors to differentiation are shown in Table 1 with their quality assessment and ranking value; the glycopeptides are listed in ranked order. None of these glycopeptides have contribution z-cores less than -2, and only one passes quality assurance metrics. Missing value induced quality issues are the primary contributors to many of the differences seen here; four of the top five ranked glycopeptides had glycans not present in the informed search space, and one (Hex 3; HexNAc 7; Fuc 5; NeuAc 2; Sulfate 1) is not bio-synthetically likely and was unobserved in the glycomic data.

**Table 1.** Glycopeptide Rankings in Naïve vs Informed AGP: YNTT

| Glycopeptide | ZScore | Quality | Likely Failure Cause |
| --- | --- | --- | --- |
| YnNT{7;6;0;1;0} | -1.947 | F | Unseen In File 2 |
| YnNT{6;5;0;3;0} | -0.934 | T | Passed |
| YnNT{7;6;1;1;0} | -0.886 | F | Low Quality In Both |
| YnNT{3;7;5;2;1} | -0.884 | F | Low Quality In Both |
| YnNT{8;6;1;3;0} | -0.880 | F | Low Quality In Both |

A truncated, reformatted ranking data summary from RAMZIS. Each row shows a different glycopeptide's identity (PEP{Hex;HexNAc;Fuc;Neu5Ac;Sulfate}), contribution z-score, quality indicator, and probable quality failure cause.

#### 3.1.2 AGP: Comparing Across Heterogeneities

RAMZIS allows for deeper understanding of the differences induced by search space differences and complexity increases. To assess the impact of complexity, we present the results of two comparisons between purified AGP (Mix 1) and Mix 5, a sample of equal parts AGP, transferrin, fetuin, haptoglobin, and alpha-2-macroglobin. The left hand side of Figure 2 pertains to a Naively searched analysis while the right hand side pertains to an Informed search analysis; all graphs in Figure 2 are in relation to AGP glycosite YNNT. Figure 2a shows the Internal similarity distributions of Mix 1 in the comparison of Mix 1 and 5. The left of 2a shows the Naïve Mix 1 to be completely disjoint with the Test similarity, leading to an IC of 3.97, an FPR and an FNR of 0%. The Informed search on the right of 2a is also disjoint with an IC of 12.18 and 0% FPR and FNR. Both Mix 1 Internal similarities pass all quality metrics, and the Test Similarity of both is simulated well by RAMZIS with observed similarities with z-scores less of 0.86 and 0.11 for Naïve and Informed respectively.

**Fig 2.**
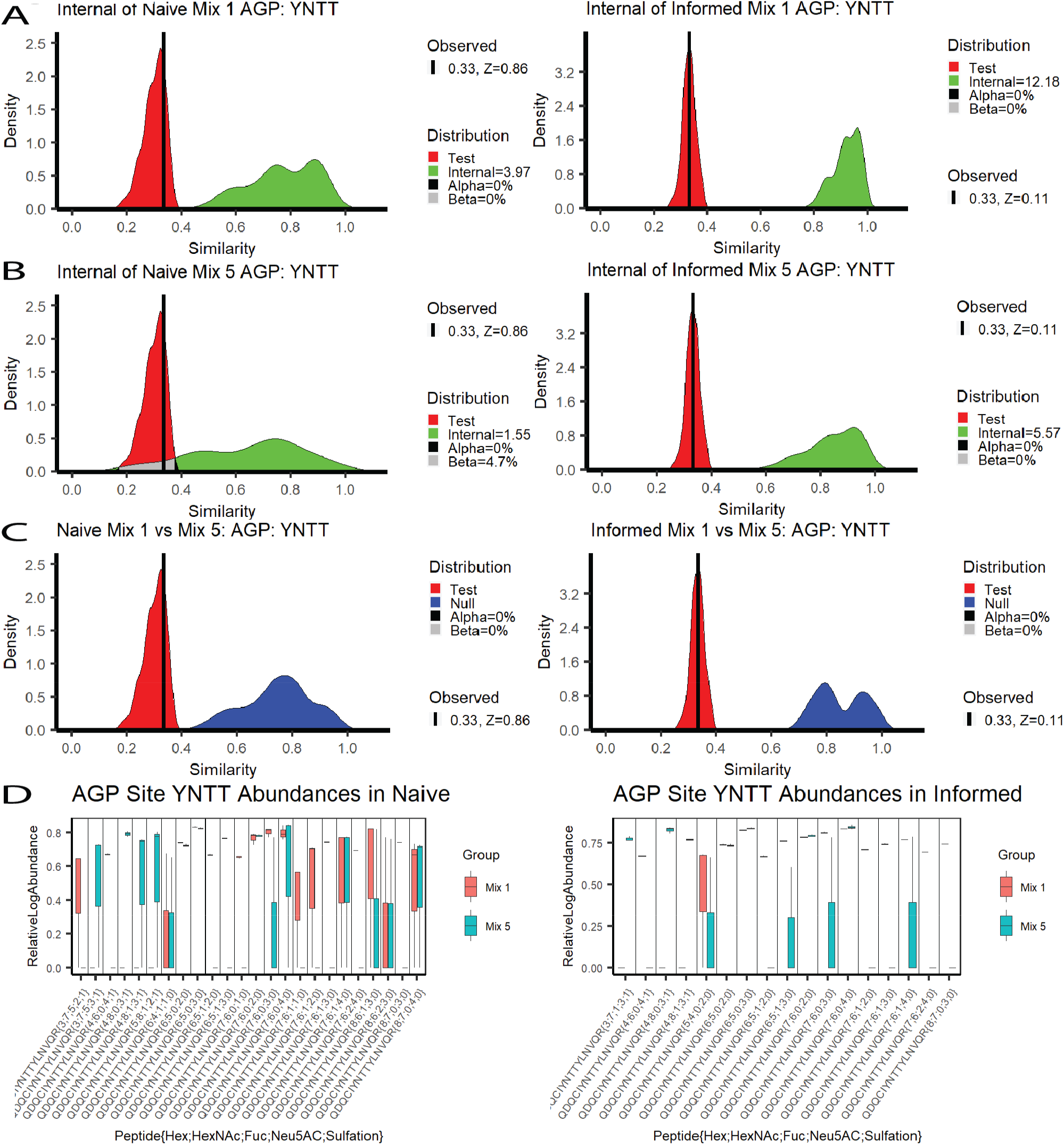
Related Similarity Comparisons in AGP Mixtures. A side-by-side display of two related similarity comparisons: purified AGP compared to AGP in a heterogeneous mixture. The left column is the naïve search while the right is the informed. A) Internal similarities of pure AGP. Naive AGP IC=3.97, 0% FPR & FNR; Informed AGP: IC=12.18, 0% FPR & FNR. B) Internal similarities of Mix 5. Naive AGP: IC=1.55, 0% FPR, 4.7% FNR; Informed AGP: IC=5.57, 0% FPR & FNR. C) Comparisons of Test and Null Distributions. The Naive comparison had a 0% FPR and a 0% FNR; the Informed comparison had a 0% FPR and a 0% FNR. RAMZIS simulated the comparison sufficiently for both identification methods with Test z-scores of 0.86 and 0.11 for the Naïve and the Informed respectively. D) The relative log abundances used to calculate the above similarities are displayed by glycopeptide in a box plot; Mix 1 is in red while Mix 5 is in green. These have been standardized by TIC and scaled according to a linear model by 1.067 and 1.084 for Naïve and Informed respectively. Mix 1 Naïve has a 70% presence rate; Mix 5 Naïve has a 39% presence rate; Mix 1 Informed has an 80% presence rate, and Informed has a 66% presence rate.

Figure 2b shows the Internal Similarity distributions of Mix 5 shows divergence in behaviors between Naïve and Informed searches. The Naïve Mix 5 on the left has an IC of 1.55, an FPR of 0%, and an FNR of 4.7%, while the Informed Mix 5 has an IC of 5.57 and an FPR and FNR of 0%. The Informed gives high enough data quality to reliably compare the two datasets, while the Naïve search space fails at the higher complexity. Subtle multimodality is present, but Mix 5 had an n=3, so outlier removal was not performed. The General Comparisons in Figure 2c both show completely disjoint Null and Test distributions for both the Naïve and Informed comparisons with 0% FNR and FPR for both.

A truncated, reformatted ranking data summary from RAMZIS. Each row shows a different glycopeptide’s identity (PEP{Hex;HexNAc;Fuc;Neu5Ac;Sulfate}), contribution z-score, quality indicator, and probable quality failure cause.

Figure 2d shows the boxplots of the TIC standardized abundances. In the Naïve, Mix 1 has 4 absent glycopeptides while Mix 5 has 10; in the Informed, Mix 1 has 3 absent glycopeptides while Mix 5 has 6. Multiplicative scaling terms were applied to both Mix 5 abundance datasets to account for a semi-uniform shift of abundances that were not reflected by the TIC standardizations. This rescaling does not substantially change the overall comparisons, and a version of Figure 2 without this shift can be found in the supplemental.

A truncated, reformatted ranking data summary from RAMZIS. Each row shows a different glycopeptide’s identity (PEP{Hex;HexNAc;Fuc;Neu5Ac;Sulfate}), contribution z-score, quality indicator, and probable quality failure cause.

Table 2 shows the ranking information for the highest ranked glycopeptides in the Naïve comparison of Mix 1 and Mix 5. All of these glycopeptides are absent from Mix 5 and have significant contributions to dissimilarity. One of the glycopeptides (Hex 4; HexNAc 6; NeuAc 4; Sulfate 1) is biosynthetically unlikely, but it was identified in the glycomic data. Examination of the GlycReSoft files shows that it’s identifying spectra share information with more biosynthetically likely glycans, but that in some cases, it was selected based off of MS1 error. A full Naïve comparison ranking summary table is available in Supplemental Table 6.

**Table 2.** GP Ranking of Naïve Mix 1 vs Mix 5: AGP: YNTT

| Glycopeptide | ZScore | Quality | Likely Failure Cause |
| --- | --- | --- | --- |
| YnTT{7;6;0;1;0} | -2.017 | F | UnseenIn_File2 |
| YnTT{6;5;1;2;0} | -2.015 | F | UnseenIn_File2 |
| YnTT{4;6;0;4;1} | -2.015 | F | UnseenIn_File2 |
| YnTT{7;6;2;3;0} | -2.008 | F | UnseenIn_File2 |
| YnTT{7;6;1;3;0} | -2.000 | F | UnseenIn_File2 |

A truncated, reformatted ranking data summary from RAMZIS. Each row shows a different glycopeptide's identity (PEP{Hex;HexNAc;Fuc;Neu5Ac;Sulfate}), contribution z-score, quality indicator, and probable quality failure cause.

Table 3 shows the top five ranked glycopeptides of the Informed Mix 1 to Mix 5 comparison. All five are unseen in Mix 5, and are all significant contributors to dissimilarity. The same unlikely glycopeptide from the Naive is seen again. A full Informed comparison ranking summary table is available in Supplemental Table 8.

**Table 3.**
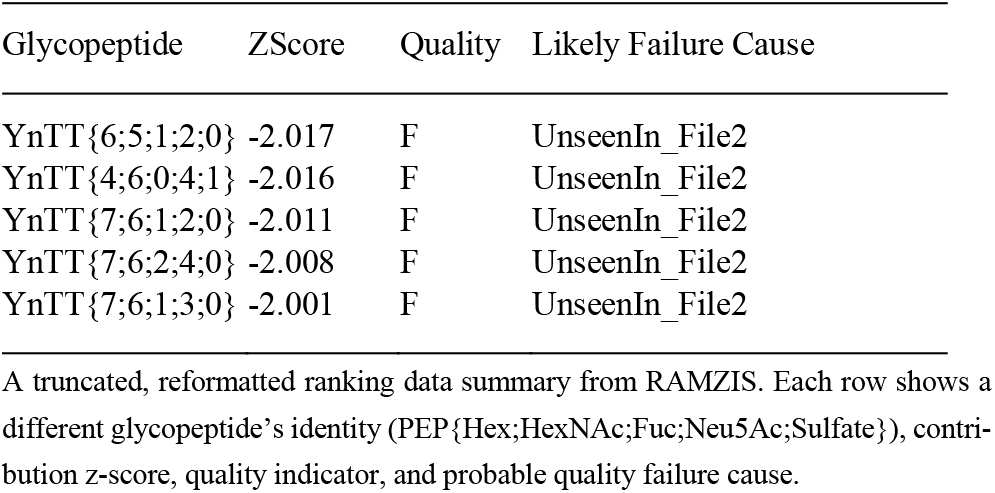
GP Ranking of Informed Mix 1 vs Mix 5: AGP: TPNK

An unscaled analysis can be found in Supplemental Figures 12 and 13 with their ranking summaries, Supplemental Tables 7 and 9 for Naïve and Informed respectively.

## 4 Discussion

### 4.1 AGP Results Interpretation

The study that produced these data showed that the use of an informed search space improved glycopeptide identification confidence. RAMZIS has confirmed and helped quantify this phenomenon.

There is an observable difference between Naïve and Informed search analysis even when just looking at purified AGP. The ranking information quantitatively confirms the visually apparent fact that this difference is not from differences in quantification, but instead from missingness caused by an overly broad naïve search space. This analysis enables a more stringent confirmation than traditional statistics allows for.

This case-study also confirms the decrease in identification confidence that comes with increased sample complexity. The Informed comparison between Mix 1 and Mix 5 shows differentiability between the two that is primarily caused by missingness in Mix 5, despite Mix 5 actually having larger glycan and protein search spaces. As mixture complexity increases, the likelihood of identifying lower abundance glycopeptides decreases. While confounded by the non-uniform scaling, there is still evidence that quantification is susceptible to this same loss in confidence. The glycopeptides that passed quality assurance metrics were still contributors to difference and were more so prior to the artificial adjustment made for the purpose of a more conservative analysis. This change in quantification confidence is not unreasonable as with less confidence in an identification there may become fewer points in the elution from which to find the area under the curve.

The Naïve comparison of Mix 1 and Mix 5 shows the two issues compounding upon one another. The naively searched Mix 5 has a much higher rate of missingness, but instead of forcing differentiation from Mix 1, it appears as a low quality subset of it. We can identify outlier issues via modality in Mix 5, and while the high degree of overlap in the general comparison should be expected of a glycosite with the same glycosylation, the degree of overlap between the Internal and Test similarities of Mix 5 show a lower degree of internal consistency than of Mix 1. There are small yet observable differences in quantification even after rescaling, and the quantifications themselves are more variable in the naïve search space than the informed as seen in Figure 2d.

One of the most important problems in glycoproteomics is determination of assignment confidence for a complex sample. We have demonstrated the use of RAMZIS to compare glycopeptide assignments and quantifications between low and high complexity glycoprotein mixtures. We found that the difference between informed and naïve searches are discernable even at low complexity, and that the effects become magnified as complexity increases. As mixture complexity increases, the identifications with higher signal values become the only ones identified, and they become less consistently quantified, especially in uninformed search spaces.

### 4.2 RAMZIS Broader Impacts

As most of the questions that glycoproteomics seeks to answer are in inherently higher complexity systems, the ability to assess data quality as part of the comparative process is key. The ability to understand not only that there are differences, but to be able to quantitatively determine the source of these differences, will be necessary for a fuller understanding in both improving our methods and interpreting our experiments.

Our data quality decreases with every additional glycopeptide that must be tracked, and we fight against technological limitations of throughput in our attempts to quantify the entire glycoproteome. We must know when our attempts to surpass these throughput problems become overwhelmingly deleterious to our ability to reliably quantify, and we must be able to identify how exactly they are so; this knowledge will enable us to fine tune methods, determine acceptable losses, and ideally, perfect them to work around those deleterious effects.

RAMZIS is an effective tool for modeling glycopeptide comparisons that gives a clear readout of assignment confidence that will guide users in appropriate interpretation of their glycoproteomics data. It is a step in the road to developing better methods and performing more consistent analysis of glycoproteomics. With it users will be able to identify outliers, assess data consistency, perform heuristic glycosite comparisons, and target potential identifications that are the sources of error or differentiation. RAMZIS is ultimately a quality assurance tool; it allows users to answer the question: “Am I able to ask the question I want to ask?” If the Internal Similarity distributions are sufficiently well defined, then it is possible for the user to ask the next question: “Are these glycosites likely to be differentiable?” And while it does enable differentiating glycosylation patterns at a glycosite, the rankings it provides are meant to steer the user towards the likely answers will inform further experimental investigations to help answer the final question: “How are these glycosites different?”

We emphasize that analysis that RAMZIS provides should be backed up with other statistical tests once the user has identified likely candidates, preferably with targeted or DIA mass spectrometry data. The most appropriate use of RAMZIS is as a guide, not a determinant. We plan to improve the functionality of the toolkit to decrease the impact of missing values. We are also testing the viability of incorporating information from the biosynthetic network into RAMZIS, which would enable a better understanding of the rankings and help to identify patterns if any glycopeptides are behaving differently than their related network neighbors.

## Acknowledgements

The authors would like to thank Margaret Downs, Meizhe Wang, and Justin Moy for assisting with user testing.

