## SupplementalTextAndFigures for "RAMZIS: a bioinformatic toolkit for rigorous assessment of the alterations to glycoprotein structure that occur during biological processes"

### RAMZIS Supplemental

This document contains elaboration on the workings of RAMZIS and presents more of the testing done.

### Supplemental Methods

#### Algorithm Workflow

Figure 1. Algorithm Workflow

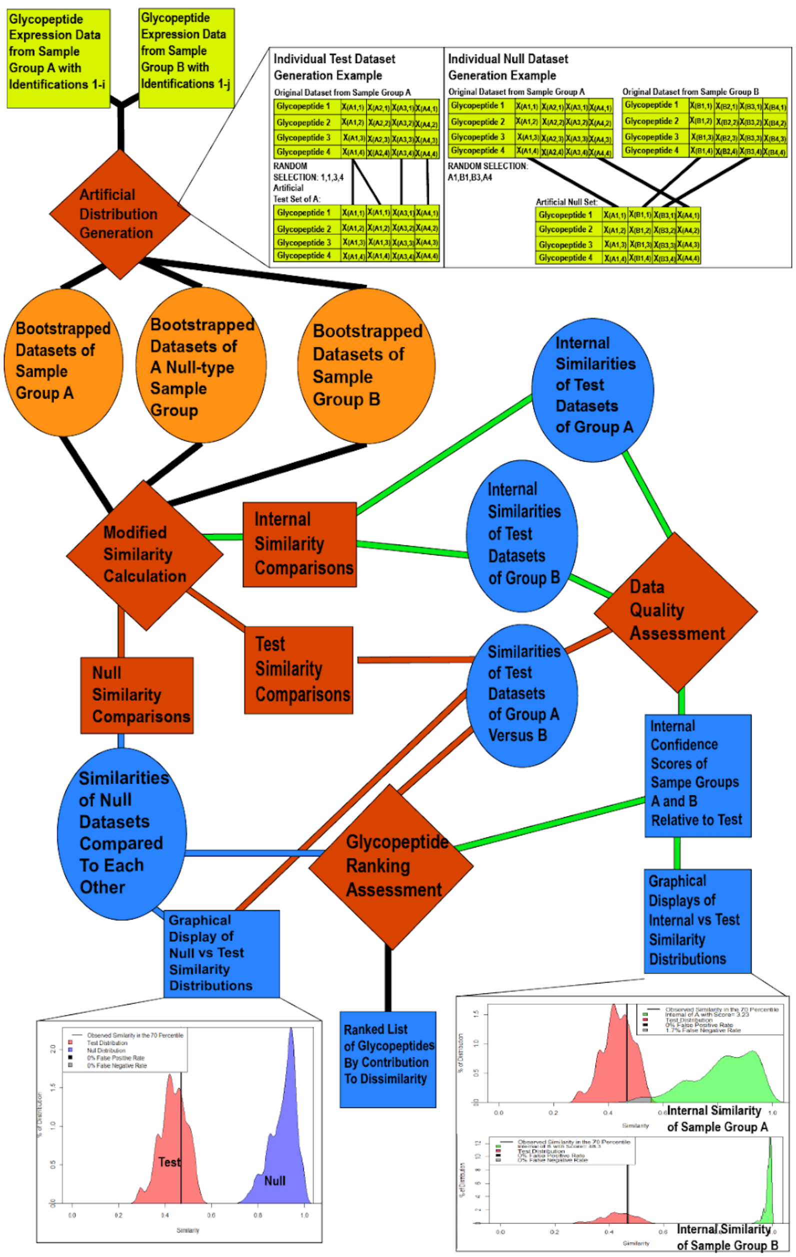

##### Supplemental Figure 1: Algorithm Workflow

Supplemental Figure 1 depicts the algorithmic workflow of RAMZIS. Inputs are depicted in yellow; processes are depicted in red; temporary data are in orange; outputs are in blue. Circles are datasets; squares are user interpretable.

- The user provides two glycopeptide expression datasets, herein called Group A and Group B
- Those datasets undergo bootstrap sampling to produce 3 different kinds of simulated datasets: Samplings of Group A, Samplings of Group B, Samplings of Groups A & B at the same time.
- These datasets are the subjected to different kinds of similarity comparisons. The Samplings of A produce Internal Similarities of Group A. The Samplings of B produce Internal Similarities of Group B. The Samplings of A compared to B produce Test Similarities. The Joint samplings produce Null similarities.
- These similarities are fed into data quality assessments and glycopeptide ranking assessments.
- The similarities and their data assessments are used to produce comparative similarity graphs.

### Input Data

RAMZIS can be used to format data from GlycReSoft outputs, but if you are using a different tool for Glycopeptide identification and quantification, you will need to do your own formatting. It's also recommended to input a normalization vector.

#### Data Format Example

The universal data format that RAMZIS accepts is a matrix with glycopeptides for rows and samples for columns. They're expected to have row and column names. Row names are how RAMZIS knows which identifications are the same between files. The cells are filled with  The matrix of one file looks something like this:

**Supplemental Table 1**

| | Sample_1 | SampleB | PatientIII | NumberD |
|---|---|---|---|---|
| GlyPep1 | 1230000 | 1320000 | 1290000 | 120000 |
| GlyPep2 | 95000 | 105000 | 100000 | 90000 |
| GlyPep3 | 20000 | 25000 | 22000 | 18000 |

RAMZIS requires two of these files as input. It will also accept dataframes formatted in this fashion.

The names of the samples (columns) don't matter so long as they don't break R formatting rules. Glycopeptides can similarly be named anything so long as they are consistent between files. It will not combine missed cleavages into a single glycosite, and any such data manipulations should be done prior to input into RAMZIS. RAMZIS can be used to compare PTMs other than glycosylation, and if other PTMs are included the glycopeptide names, they will be treated as fully separate glycopeptides from equivalent glycopeptides without those PTMs as RAMZIS uses an exact match system.

#### Standardization & Relativization

There are multiple standardization methods involved in the initial data processing.

**Log Transformation**

Users may choose to not have their data undergo log transformation(logopt=F), but it is recommended(Default: logopt=T). RAMZIS uses the default log() function of R, which is the natural log. Log transformation of most glycopeptide abundance data scales the overall data such that it becomes more normally distributed.

**Minimum Observation Requirements**

For a glycopeptide to be considered real it must be seen twice(Default: kmin=2) between the two datasets. This requirement for cross dataset minimums is based on the idea that the two datasets, until proven otherwise, are from the same underlying glycosylation distribution. The number of observations required to be seen may be changed, and the cross dataset minimum may be disabled. Internal Similarity is calculated without a minimum observation requirement to make comparisons more conservative; low observation identifications reduce the overall similarity and make overlap between the Test and Internal more likely. Users may also set the internal minimum observation threshold. (Default: kmin_int=1)

**Normalization aka Standardization**

Data is not normalized in RAMZIS, but it can be standardized. Normalization, the process of turning values into z-scores, is avoided as the similarity metric requires positive values. Standardization, the process of scaling abundance values between samples and datasets, is optional but recommended. (Default: normvec=list("None","None")

RAMZIS expects two vectors in a list for its standardization process, wherein the first vector should correspond in length to the number of samples in the first input dataset and the second vector should correspond in length to the number of samples in the second input dataset. The first sample in the first dataset will have all its glycopeptides multiplied by the first value of the first vector in the list, the second sample in the first dataset will be multiplied by the second value of the first vector and so on.

The source of these standardization factors is up to the user, but it is recommended to use a factor derived from the TIC- the Total Ion Chromatogram, a single summation of the entire signal produced by the elution. If using the default log transformation, take the log of each sample's TIC and then take the inverse of it. (scaling_factor[1]=1/log(TIC[1]) If not using the log transform then simply take the inverse of the TIC. (scaling_factor_nolog[1]=1/TIC[1])

Another potential source of standardization is from protein abundance data. If a protein is more abundant in one sample group over another, then the TIC may not properly adjust. In this case, the use of a more complex scaling term would be appropriate. This could take the form of a series of conversion factors like this:

$$ScalingFactor(Sample_i, Protein_j) = \frac{1}{ln(TIC_i)} * \frac{E[Abundance_j]}{Abundance_{j,i}}$$

Where i is a sample index, and j is a given protein. E[Abundance(j)] would be the mean abundance of the protein in all samples across datasets. Abundance(j,i) would be the abundance of the protein j in the proteomics data that corresponds to sample i.

**Joint Relativization**

Joint Relativization is performed when no standardization vectors are given; it can be enabled to occur even then (Default: rel_force=F). Relativization divides the abundances given by It defaults to making all abundances relative to the largest summed abundance of any sample in both datasets. (Default: rel="Joint") This default assumes that the overall signal is equivalent between files, and that differences between the files can be believed to be from concentration rather than signal response variance. There are also options to make abundances relative to the largest summed abundance within a dataset (rel="Within"), to be left as is (rel="AsIs"), to have each sample relative to its own total abundance (rel="Self"), or have all values relative to a specified value (rel=NUMERIC).

### Data Simulation via Bootstrap

**Test Dataset Generation**

Each input dataset will have simulated data generated from it. A simulated dataset will be a random sampling by column with replacement. 100 samplings are produced, and in lower sample number projects the samplings will be combinatorially exhaustive, but will not contain repeats and thus have fewer than 100 samplings. These Test datasets will be used to produce the Internal Similarity Distributions, by comparing all those of a given origin dataset to all others in the origin dataset, and the Test Similarity Distribution, by comparing all samplings of one dataset to all samplings of the other dataset.

**Null Dataset Generation**

To generate the two Null Datasets required, random samplings of a joint dataset are taken with probability equally split between sample groups, and probability split within those groups equally to each sample. There is a minimum number of samples required from each dataset so that there are no full exclusions in a given sampling. Repeated samplings are not allowed. If the origin datasets have equivalent sample size then the samplings will be produced together and then split evenly to produce two Null samplings. If the origin datasets have unequal sample sizes then they are produced separately.

### Similarity Comparison

The similarity equation is largely explained in the article itself. This section is to denote two features that can be modified within the analysis.

**Missing Value Correction**

This is not an imputation option. As a default, RAMZIS fills in unseen glycopeptides with zeros. (Default MVCorrection=T) If this option is disabled (MVCorrection=F) then the P terms of the similarity equation are disabled, meaning that missingness will no longer factor into similarity. This substantially changes similarity comparisons with high degrees of missingness and may induce errors in cases of complete absence.

**Distance Scaling**

Distance scaling (K) can be changed. (Default: mn=F) As a default, it has K=1+mean(Presence(A),Presence(B)) such that K can be at most 2, meaning that in cases of maximal distance the numerator will be halved. If this option is given a numeric value, it will use that value. This is convenient if the user wishes to increase the impact of the manhattan distance between glycopeptide abundances.

### False Positive and False Negative Rates

Equations 3 and 4 are presented in the paper are simplified with T as the density of the Test Distribution; R as the density of a Reference distribution; x as the point at which Test has a lower density than the Reference. These densities are approximated by the empirical cumulative distribution as determined by R’s ecdf() function.

$$AlphaOverlap(T,R) \ = \frac{\int_x^1 T}{T+R-(\int_x^1 T+\int_0^x R)} \qquad (3)$$

$$BetaOverlap(T,R) \ = \frac{\int_0^x R}{T+R-(\int_x^1 T+\int_0^x R)} \qquad (4)$$

Are simplified forms as there can be multiple inflection points wherein the reference and test distributions cross over one another. Inflection points are determined using R’s stats::density() function with set bounds from -0.1 to 1.1; once generated these densities are scaled to their totals. More accurate versions of equations 3 and 4 can be represented here:

$$AlphaOverlap(T,R) \ = \frac{\sum_i^{(n)}(ecdf(T)[i[2]]-ecdf(T)[i[1]])}{T+R-(\sum_i^{(n)}(ecdf(T)[i[2]]-ecdf(T)[i[1]])+\sum_j^{(m)}(ecdf(R)[j[2]]-ecdf(R)[j[1]]))} \qquad (3)$$

$$BetaOverlap(T,R) \ = \frac{\sum_j^{(m)}(ecdf(R)[j[2]]-ecdf(R)[j[1]])}{T+R-(\sum_i^{(n)}(ecdf(T)[i[2]]-ecdf(T)[i[2]])+\sum_j^{(m)}(ecdf(R)[j[2]]-ecdf(R)[j[1]]))} \qquad (4)$$

Here, n represents pairs of similarities, i, where i[1] is where the Test density becomes lower than the Reference and i[2] is the last similarity that the Test is lower than the Reference; m represents pairs of similarities, j, where j[1] is where the Reference density becomes lower than the Test and j[2] is the last similarity where the Reference is lower than the Test. No region is double counted.

### Internal Confidence Score

**Weighted Z-Score**

The current Internal Confidence Score follows equation 5:

$$InternalScore \ = \ (\frac{\mu_{Test}-\mu_{Internal}}{\sigma_{Internal}}) \ * \ 10^{-(\alpha+\beta)}$$

Where μ are means of their respective similarity distributions; σ is the standard deviation; α is the false positive rate from equation 3, and β is the false negative rate from equation 4. This weighted z-score is penalized by larger overlaps to avoid overconfidence in case of long tailed distributions, which are more common in non-log transformed similarities. Joint overlaps greater than the combined 0.25 (from 0.05 and 0.20 for FPR and FNR respectively) will more than halve the z-score.

**Prior Version Scoring: Pseudo-Kurtosis**

A prior version of the Confidence score used the similarity density height to variance as a form of kurtosis like measurement. That used a threshold of Score>100, meaning that an Internal Similarity Distribution would have a density plot height 100 times greater than the similarity variance. This was heavily reliant on density plotting, not necessarily user interpretable, and not as well grounded in mathematics.

### Ranking Identifications

The ranking and quality assessment of individual glycopeptides contributions to similarity rely heavily on two different related measurements. That is the unweighted contribution to similarity, and the weighted contribution.

$$UnweightedContribution(g) = A_g B_g K^{-d(A_g, B_g)}$$

$$WeightedContribution(g) = \frac{A_g B_g K^{-d(A_g, B_g)}}{\sum_i^G (A_i^2) + \sum_i^G (B_i^2) - \sum_i^G (A_i B_i K^{-d(A_i, B_i)})}$$

The unweighted contribution is just the numerator of the modified similarity equation while the weighted contribution is adjusted by the denominator term from the similarity equation. The former is used in ranking, while the latter is used in quality assessment.

#### Quality Metrics

As the contributions have much smaller distributions, quality thresholds for them are joined from an alpha of 0.05 and beta of 0.20 to a joint overlap (of Internal and Test) proportion of 0.25. The Internal Similarity contribution should be greater than the Test if using the Weighted Contributions. Other quality metrics have been identified, but are not currently implemented into the main workflow as they are still being tested.

**Weighted Contributions**

The weighted contributions are used for quality assessment as they provide a more conservative measure of overlap between the Internal and Test distributions. If a user desires for it to be consistent with the ranking process, they may specify to the less conservative unweighted contributions. (QualityInfo=”Numerator”)

#### Ranking by Contribution

The ranking of an individual glycopeptide uses the average z-score of its Test similarity contribution in relation to the Null Similarity contribution distribution.

$$RankingInfo = \frac{1}{n}\sum_i^n \left(\frac{Contribution_{Test,i} - \mu_{Contribution,Null}}{\sigma_{Contribution,Null}}\right)$$

Where i is one comparison of all n Test similarity comparisons. The lower the z-score, the more likely it is that the glycopeptide in question contributes strongly to the differences in the overall similarity distribution.

**Unweighted Contribution**

While the unweighted contributions are less conservative when used in the quality assessment step, the weighted contributions are misleading when used as the ranking information. Missing values are the cause of this phenomenon. When there is a complete absence of a glycopeptide in one dataset, then the Test contributions for that glycopeptide will all be zero, but the Null similarity contributions will have high variance, and may include zeros. This produces a high variability in the Null, leading to a lower absolute z-score for that glycopeptide, which isn't problematic, highly variant glycopeptides should not be considered consistent contributors to dissimilarity. But other glycopeptides will have higher weighted contributions if there are low Null values boosting their value.

Use of weighted contributions for ranking information causes missing values to be devalued in rankings. Weighted contributions are biased by the effect of missing value glycopeptides on fully present glycopeptides, and so using weighted contributions can lead to spurious conclusions. Unweighted contribution based ranking, while less conservative, acknowledges the impact of missing values and does not boost other glycopeptides.

Should a user wish to use weighted contributions they may do so, but it is not recommended. (RankingInfo="WeightedContributions")

**Alternative Quality Metric When Using Weighted Contributions**

If using the weighted contributions for ranking, a secondary confirmation on a significant glycopeptide ranking can be helpful to be sure that there is no bias from missing values, however this is not a guaranteed filter. This second confirmation is a comparison of the normalized contribution of the Test against the normalized contribution of the Null. For a given glycopeptide, generate the z-score of the contribution compared to all other contributions in the similarity distribution in a leave-one out fashion.

$$Z(g_{SimDis}|G_{SimDis}) = \frac{g_{SimDis} - \mu_{G!=g}}{\sigma_{G!=g}}$$

Where $g_{SimDis}$ is the similarity contribution of a given glycopeptide in a Similarity Distribution SimDis; $G_{SimDis}$ are all contributions of all glycopeptides in SimDis; $\mu_{G!=g}$ is the mean of all contributions in G excluding the given glycopeptide g, and $\sigma_{G!=g}$ is the standard deviation of all contributions in G excluding the given glycopeptide g.

This z-score is generated for the Test and the Null for each glycopeptide, and only glycopeptides that have Test z-scores lower than their Null z-scores can be considered as a potentially significant contributor to dissimilarity.

### Theoretical Data Generation

The theoretical data generator is good for testing potential behaviors in RAMZIS. It allows a user to produce RAMZIS formatted datasets to their specifications. The user controls the numbers of samples and glycopeptides, as well as their variability, missingness, and their means.

**Data Generation via Beta Distribution Sampling**

The data points are sampled from a beta distribution. Beta distributions have means defined by the ratio of an alpha parameter to a beta parameter. By inputting a list of alphas and betas equal to the number of glycopeptides, the user can define the mean for each theoretical glycopeptide. Variance of a beta distribution is proportional to the size of the alpha and beta, the greater they are the less variant they are. Use of this property allows users to specify how consistent their data is separately from their alpha and beta inputs, and it can even be controlled on a per glycopeptide level. The data generation uses three variables to control the variance: 1) the number of significant figures kept of the initial mean samplings. (Default=2) This number is turned into a ratio of alpha and beta to replicate the mean in sampling; the more numbers kept after the decimal, the larger the numbers in the ratio, the smaller the variance. 2) A ratio multiplier (Default=10) which scales the numbers in the ratio, and 3) a multiplier exponent (Default=1) to more quickly change the variance by exponentiating the multiplier.

**Missingness Generation**

After data generation via beta distribution, missingness is introduced to the dataset. Based off of an average presence input by the user, each glycopeptide has its probability adjusted by abundance using the following formula:

$$P_{adjust,g} = P_{input} * e^{mean(abundance_g) - mean(abundance_{all})}$$

Wherein a glycopeptide, g, has a probability equal to the input probability multiplied by e raised to the power of the mean of the abundance of g less the mean of all glycopeptides in the dataset. Any glycopeptides that are adjusted to a probability greater than 1 are set to 0.99999999, and any that are adjusted to less than 0 are set to 0.00000001. If presence is set to 1, then the probability is not adjusted for any glycopeptide. This adjusted probability is then used in an binomial distribution to determine the number of samples that keep their abundances for that glycopeptide. The glycopeptide then has a random number of its remaining -if any- glycopeptides set to 0.

### Outlier Detection

**Modality Membership Proportion**

Outlier Detection uses the Internal Similarity distributions and their modality. If an Internal Similarity Distribution is multimodal, peak membership can be determined by assessing the samplings that produced similarities between the troughs of each given peak. A leave one out z-score assessment is then used to determine over or under-representation in the peak with a cut off of greater than absolute z-score. Samples that are under-represented in the primary (highest similarity) peak or over-represented in secondary peaks are considered deleterious outliers and should be manually removed from the dataset by the user. Peaks that are over-represented in the primary peak or under-represented in secondary peaks are not deleterious outliers as they show higher consistency than the average sample.

### Supplemental Results

#### Theoretical Examples

This section serves to show first how the theoretical data generator works. Then show different examples of comparisons. These figures are aimed at both proving validity and demonstrating use.

##### Supplemental Figure 2: Theoretical Data Generation

This graph shows the impacts of some of the modifications. The data in these plots are produced with default settings except for the means, the number of significant digits for the means, and the presence. The greater the number of significant digits, the less the variance becomes. Presence is also much more heavily affected in lower abundance values, which aligns with general expectations. The means were determined by 2 input vectors that are reverses of one another: 4E4,3.6E4,3.2E4,2.8E4,2.4E4,2E4,1.6E4,1.2E4,8E3,4E3. Generating the means: 0.0476…,0.142…,0.238…,0.333…,0.428…,0.523…,0.619…,0.714…,0.809…, and 0.904…

**Figure S2: Theoretical Data: Variance vs Presence**

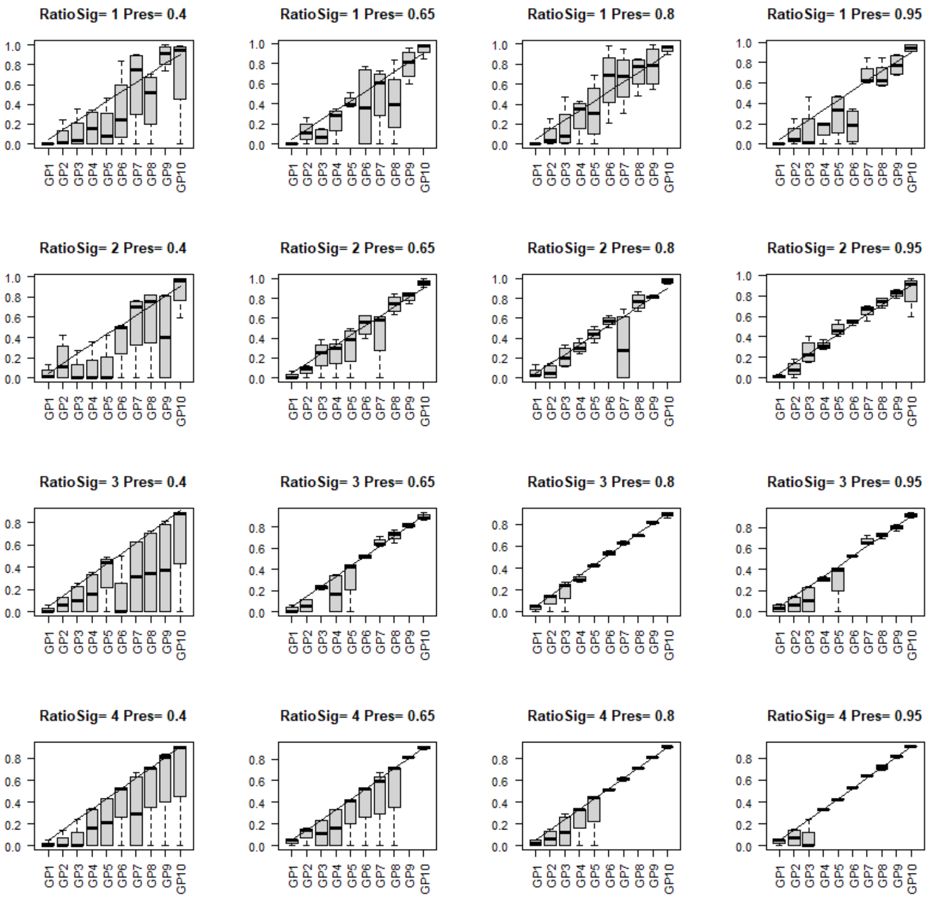

#### Theoretical Case: Same Underlying Distribution

Before proving that RAMZIS's similarity comparisons can find differences between groups, we must first prove that it finds groups with the same underlying glycosylation patterns to be similar.

For this, we generated a 12 glycopeptide dataset with 10 samples, low variance conditions (significant digits of mean=2, ratio multiplier=10^2 (Default=10^1)), and a presence of 1 with sampling
means=c(0.0706,0.152,0.229,0.303,0.381,0.454,0.538,0.617,0.703,0.762,0.842,0.926) from initial alpha=c(400,800,1200,1600,2000,2400,2800,3200,3600,4000,4400,4800) and initial beta=c(4800,4400,4000,3600,3200,2800,2400,2000,1600,1200,800,400).

Then those 10 samples were split into two different datasets. This produced these abundances and similarity comparisons.

**Supplemental Figure 3: Theoretical: Same Underlying**

**Supplemental Figure 3: Theoretical: Same Underlying**

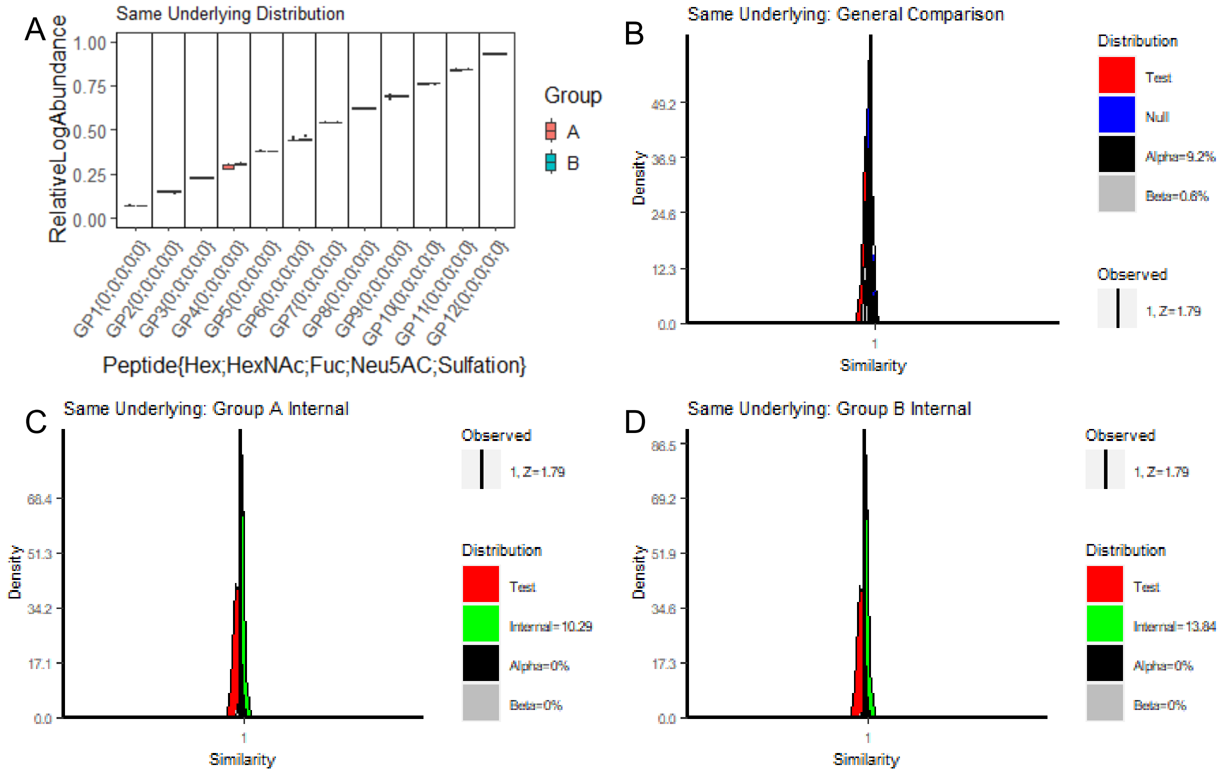

The abundances in S3a show very little variance between the groups. Figure S3b is the general comparison (between the Test and Null), which shows an FPR of 9.2% and FNR of 0.6%, exceeding the FPR threshold of 5%, so the two groups cannot be differentiated. The simulation of the comparison is within reliable ranges with an absolute z-score less than 2 at the Observed Similarity equal to 1 after rounding. We can say this confidently as seen in S3c and S3d where the Internal similarities are well over the confidence threshold and there is no substantial overlap between them and the Test similarity distribution. With this example, we can confirm that RAMZIS won't find a spurious difference in the ideal circumstance. Note that the similarity plots displayed in S3 and S4 are shown with the x-axis confined to 0.9 to 1.1 for easier viewing.

**RAMZIS Simulation Failure**

To better resemble the abundances and variances seen in the AGP data in Figures 1a & 2d, we created another dataset with a decreased ratio exponent (10^1) compared to the prior dataset (10^2) to increase variability, and an initial alpha=c(1700,3550,3000,3660,6280,3720,3560,3920,3980,4480,6112,5000) and an initial beta of 2000 repeated 12 times; this produced sampling means=c(0.081,0.162,0.197,0.202,0.23,0.258,0.282,0.327,0.391,0.465,0.547,0.742) & expected variances=c(7.4e-06,2.7e-05,1.6e-05,3.2e-05,1.8e-4,3.8e-05,4e-05,2.2e-05,2.4e-05,1.2e-04,2.5e-05,3.8e-05) respectively.

##### Supplemental Figure 4: Theoretical: Same Underlying Increased Variance

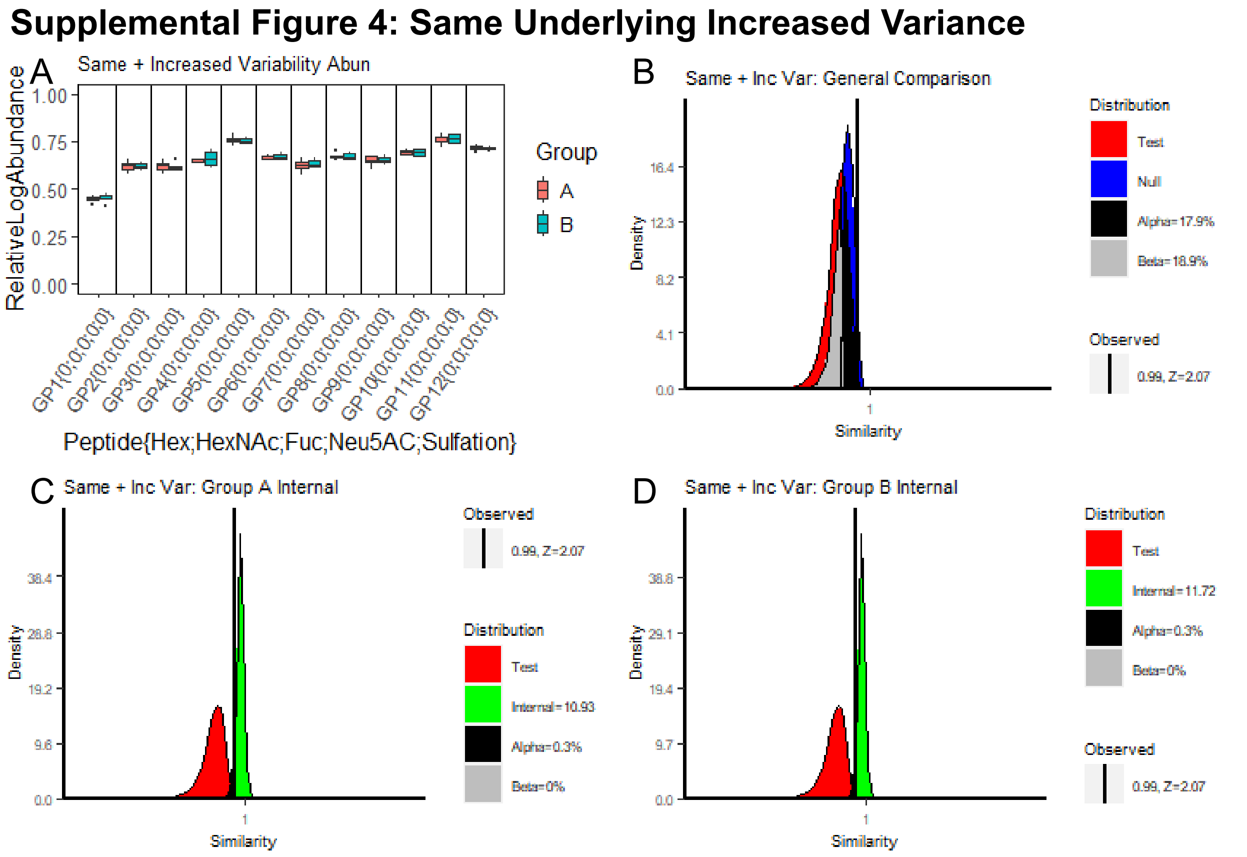

S4c and S4d show the Internal similarities of the two groups, and they both have high, tightly defined similarity distributions with high confidence scores. S4b shows the general comparison between the Test and Null with an FPR of 17.9% and an FNR of 18.9%. The Test distribution has the Observed similarity beyond acceptable bounds at 0.99 with a z-score of 2.07. This failure of simulation is increasingly likely in highly similar distributions as there is only so close to 1 that a similarity score can become. Less conservative approaches could put the z-score threshold at an absolute value of 3, but that would be an even less confident heuristic. While RAMZIS is not falsely differentiating datasets from the same underlying distribution here, it is ultimately because it cannot reliably simulate the comparison of two datasets that are so highly correlated, not because of the Test and Null overlap.

In fact, if we were to run Group A against itself in RAMZIS, as in Supplemental Figure 5, we see that RAMZIS fails once more in simulation as the observed similarity of the initial comparison is 1 as they are the exact same dataset.

#### Supplemental Figure 5: Theoretical: Group A vs Itself

**Supplemental Figure 5: A vs A Comparison**

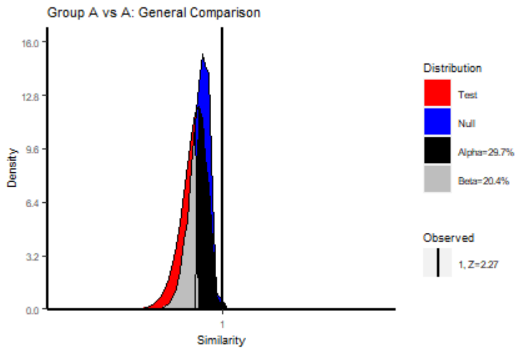

This simulation failure is a second fail safe among other quality checks. In highly consistent, highly similar datasets it can be easy to find small variations. If a dataset is well defined, then any variance from individual samples could falsely delineate the comparison. But if the samples are not outliers but are just not representative of specific glycopeptides, they will push the simulated Test comparisons to be lower similarities, so low that the Observed similarity can’t be considered within its distribution. These glycopeptides can even be identified by the ranking information.

**Ranking Elucidates Simulation Failure Causes**

Supplementary Table 2 contains the Ranking Summary Table of the comparison found in Supplemental Figure 4, wherein we compared the more variable glycopeptides from the same underlying distribution.

#### Supplemental Table 2: Theoretical Rankings of Same + Inc

| Glycopeptides | ZScore | PassOverall | FailCause |
|---|---|---|---|
| GP6 | -0.32466 | FALSE | LowQualityIn_Both |
| GP12 | -0.23901 | FALSE | LowQualityIn_Both |
| GP7 | -0.22452 | FALSE | LowQualityIn_Both |
| GP5 | -0.21336 | FALSE | LowQualityIn_Both |
| GP9 | -0.20248 | FALSE | LowQualityIn_Both |
| GP2 | -0.18938 | FALSE | LowQualityIn_Both |
| GP1 | -0.18508 | FALSE | LowQualityIn_Both |
| GP8 | -0.12249 | FALSE | LowQualityIn_Both |
| GP11 | -0.09622 | FALSE | LowQualityIn_Both |
| GP3 | -0.08148 | FALSE | LowQualityIn_Both |
| GP10 | -0.07377 | FALSE | LowQualityIn_Both |
| GP4 | -0.05271 | FALSE | LowQualityIn_Both |

No glycopeptide is significantly separated here, but their rankings give us insight into the source of the Test similarities failure as a simulation. The two highest ranked glycopeptides are GP8 and GP2 can be seen to have visible outliers and wide variances respectively in Figure S4a. They both fail quality metrics due to low quality; GP8 fails only in B, as it has a wider variance than in A, while GP2 fails in both datasets due to their variance relative to the comparison.

The next 3 glycopeptides all pass Internal quality metrics, which may feel strange visually, as GP4 in B is seemingly the most variant glycopeptide quantitation in the entire dataset in S4a. What is unseen is that within that variance there are clusters. GP4 in B has the sample abundances 0.62, 0.61, 0.70, 0.71, and 0.66. Compared to GP2 in B with abundances of 0.63, 0.61, 0.58, 0.66, and 0.63, it is easy to identify that GP2 has a more uniform spread in its abundances, even if they are over a smaller area. The groupings of GP4, being farther from the overall average of GP4, will produce weighted Test similarity contributions that are substantially lower than those in its Internal Similarity as there will be other samplings of itself that aren't as far in average abundance to compare against. This separation provides the clarity that the Internal and Test behaviors are not the same, which allows the comparison of the Test against the Null. This presents a case wherein standard assessments of variability may not represent glycopeptide behaviors well as there is an assumption of sampling size that cannot be met in glycoproteomic experiments.

#### Theoretical Case: Completely Different

As a briefer demonstration of proof of concept, the comparison shown here is from two theoretical glycosylation patterns that are completely different. Supplemental Figure 6 shows the boxplot abundances in the top-right, the general comparison in the bottom left, and the internal similarities in the remaining corners. Group A has means=c(0.077,0.154,0.231,0.308,0.385,0.462,0.538,0.615,0.692,0.769,0.846,0.923) and group B has the same means but in reverse order; additionally group A has five samples while group B has ten. The glycopeptides are named GP1 to GP12 from left to right with the above corresponding means. RAMZIS can easily simulate this comparison as shown by the z-score of the observed in the Test of 0.11. The Internal similarities are both highly separated from the Test similarity with ICs of 114 and 404, and the Null is well separated from the Test with 0% overlap and thus 0% FNR and FPR.

The multimodality observed in the Null stems from the equal probability sampling of the two differently sized sample groups; while groups are equally likely to be observed, individual samples from the smaller group appear more frequently than individual samples from the larger group. This, in addition to the disparate means produced by these samplings, make multimodal effects in Null comparisons more likely to appear. While the effect size of this phenomenon is not easily quantifiable, it is a qualitative data point that indicates the two sample groups do not share an underlying distribution.

Supplementary Table 3 holds the rankings for this comparison of completely different glycosylation patterns. The rankings here show that the rankings align with our intuitive understanding of the differences in this dataset. The highest ranked glycopeptides are those with the biggest differences between their means, GP1 and GP12. This trend holds true as the rankings descend from outermost to innermost, where there is the least difference between theoretical means. The quality assurance metrics show the conservative nature of these checks as seen in GP4 where the increase of variation was enough to identify it as a potential uncertainty, and in a real experiment, GP4 could be targeted in follow up experiments to discern its behavior if needed. Between GP4 and GP11, no glycosylation passes the z-score threshold for significant contributions to differences in similarity, further emphasizing the conservative nature of these recommendations. The z-score data is presented to the user for them to make their own assessment of the comparisons as they may wish to investigate edge cases like GPs 10 and 9, which both have between 2-3% chance of their mean coming from the Null contributions.

#### Supplemental Table 3: Theoretical Rankings of Completely Different

| Glycopeptides | ZScore | PassOverall | FailCause |
|---|---|---|---|
| GP12 | -2.612433122 | TRUE | Passed |
| GP1 | -2.581082019 | TRUE | Passed |
| GP11 | -2.529310851 | TRUE | Passed |
| GP2 | -2.471415015 | TRUE | Passed |
| GP3 | -2.370663323 | TRUE | Passed |
| GP4 | -2.240008857 | FALSE | LowQualityIn_File1 |
| GP10 | -1.973772576 | FALSE | LowQualityIn_File2 |
| GP9 | -1.958962841 | TRUE | Passed |
| GP8 | -1.875148407 | TRUE | Passed |
| GP5 | -1.663501443 | FALSE | LowQualityIn_File1 |
| GP6 | -0.7712896788 | TRUE | Passed |
| GP7 | -0.5313232507 | TRUE | Passed |

**Supplemental Figure 6: Theoretical: Completely Different**

**Supplemental Figure 6: Theoretical Opposites**

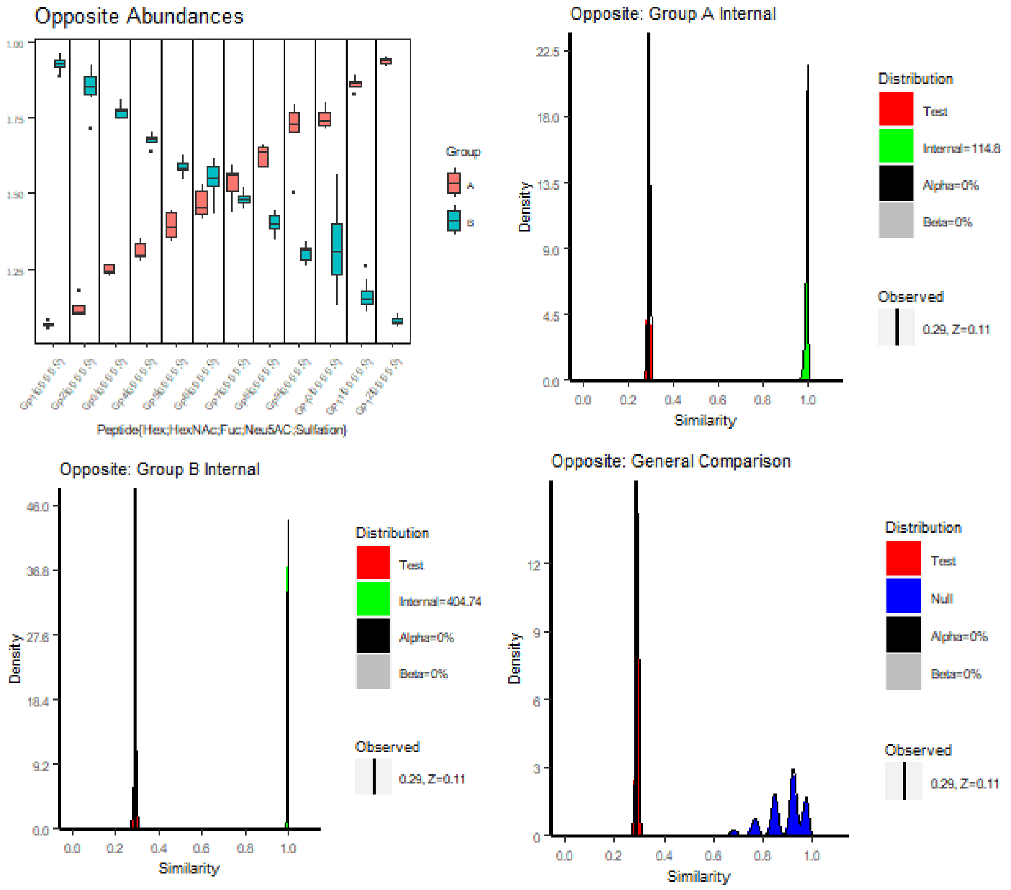

#### Theoretical Case: Major vs Minor Deviations

This section serves to show the effects of major and minor deviations on similarity comparisons. The major deviation is meant to show that RAMZIS does not wash out large individual shifts. The minor deviations section shows how small changes can lead to an overall shift in similarity.

##### Major Deviation

Using the same data as the Same Underlying example from Supplemental Figure 3, Supplemental Figure 7 shows the impact of a major deviation to one of the glycopeptides, GP9, wherein group B's GP9 has had its relative log abundance halved, which would indicate a fold change in relative log space and an even greater change in relative abundance space. All quality checks are passed; the Internal similarities are well separated and the observed is well simulated. There is no overlap between the Test and Null comparisons, and the Null is showing evidence of multimodality.

#### Supplemental Figure 7: Theoretical: Major Deviation

**Supplemental Figure 7: Major Deviation**

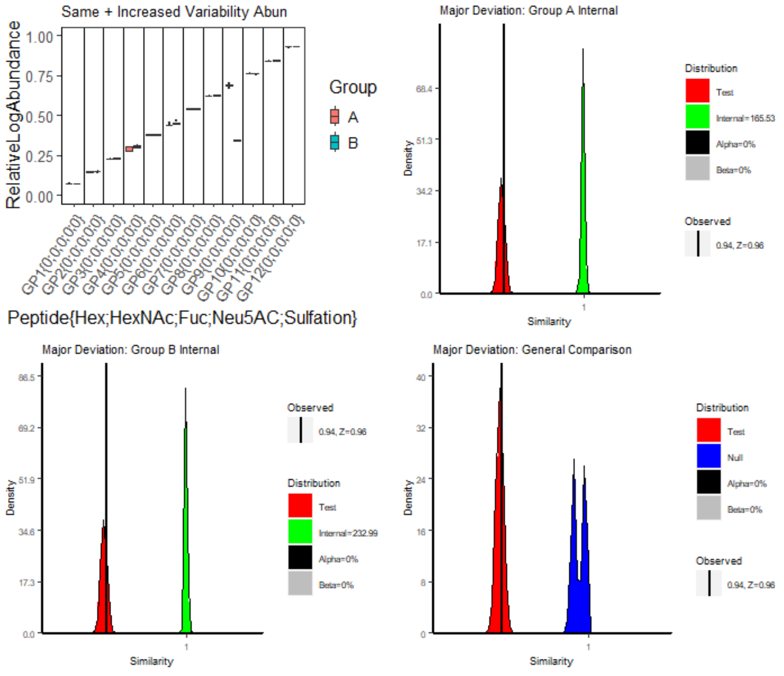

That presents the ideal case for a major deviation from an individual, a tightly defined glycopeptide undergoing a major change while remaining tightly defined, but that isn't representative of expected performance. The next example, Supplemental Figure 8, shows a major shift when variance is not as tightly defined; this data was generated by multiplying the tightly defined samples by different scalars from a normal distribution with mean of 1 and standard deviation of 0.05. The variability found here much better reflects that seen in real world cases. The left hand side shows the increased variance without the GP9 deviation, while the right has the comparison after reducing GP9 in group B by 20%; these abundances can be seen in S8a.

#### Supplemental Figure 8: Theoretical: Major Deviation in Expected Variance

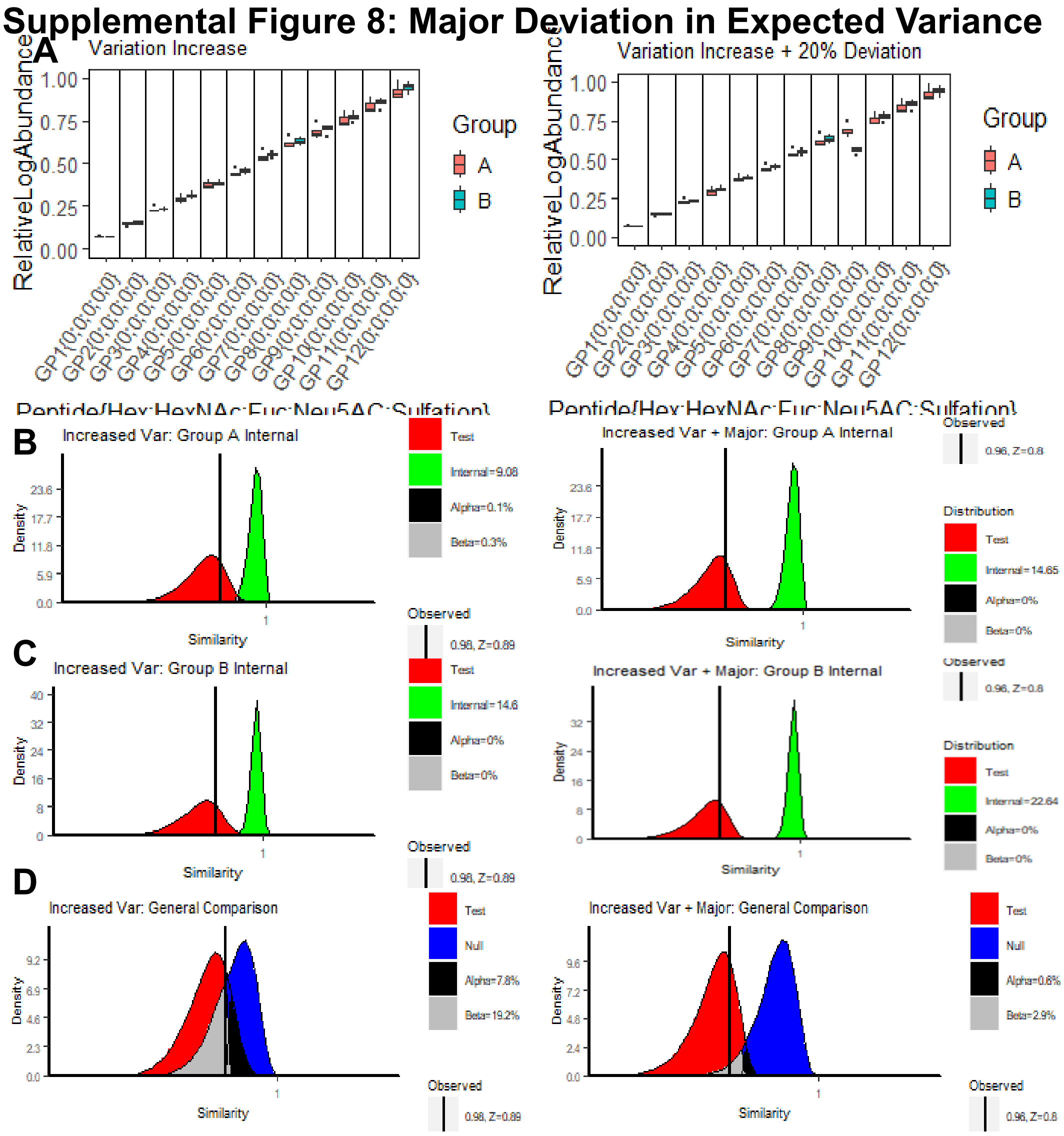

S8b and S8c show the Internal similarities which are well separated from the test in both scenarios with the same underlying having ICs of 9.08 and 14.6 while the deviation added scenario on the right has ICs of 14.65 and 22.64. Both Test similarities are well simulated with z-scores of 0.89 and 0.8 respectively, but while the left-hand scenario has too high an FPR (7.8%) to be differentiable, the right-hand (deviation added) scenario has an FPR of 0.6% and an FNR of 2.9%, allowing us to identify the difference in glycosylation pattern. The ranking for these are as expected with GP9 as the highest ranked in the deviation scenario with a z-score of -2.02 and a passing quality check; no other identifications passed quality or significance thresholds in either scenario. The 20% reduction for GP9 is actually near the edge of its detectable effect size; below this point, the glycosylation pattern change may be discernable, but GP9 no longer passes the -2 z-score threshold. 20% is not the minimum effect size for all identifications in this dataset, just the minimum for GP9. Minimum effect size is dependent on abundance magnitude and variation. As variation increases so too does minimum effect size. As magnitude increases, minimum effect size decreases; our highest abundance identifications tend to be our most confident, and our least abundant tend to be our least confident identifications. This phenomenon makes it so that higher effect sizes are required to differentiate low confidence identifications.

##### Supplemental Figure 9: Minor Deviations: Overall changes without Individual significance

Minor deviations in this section refer to deviations that would each be under their minimal effect size normally. For the example in Supplemental Figure 9, the increased variance dataset from Supplemental Figure 8 was used, but in group B: GP9 was reduced by 15%, GP3 was reduced by 20%, and GP7 was increased by 10%. These shifts can be observed in S9a. None of these pass their minimum effect size for RAMZIS.

**Supplemental Figure 9: Minor Deviations Lead to Global Change**

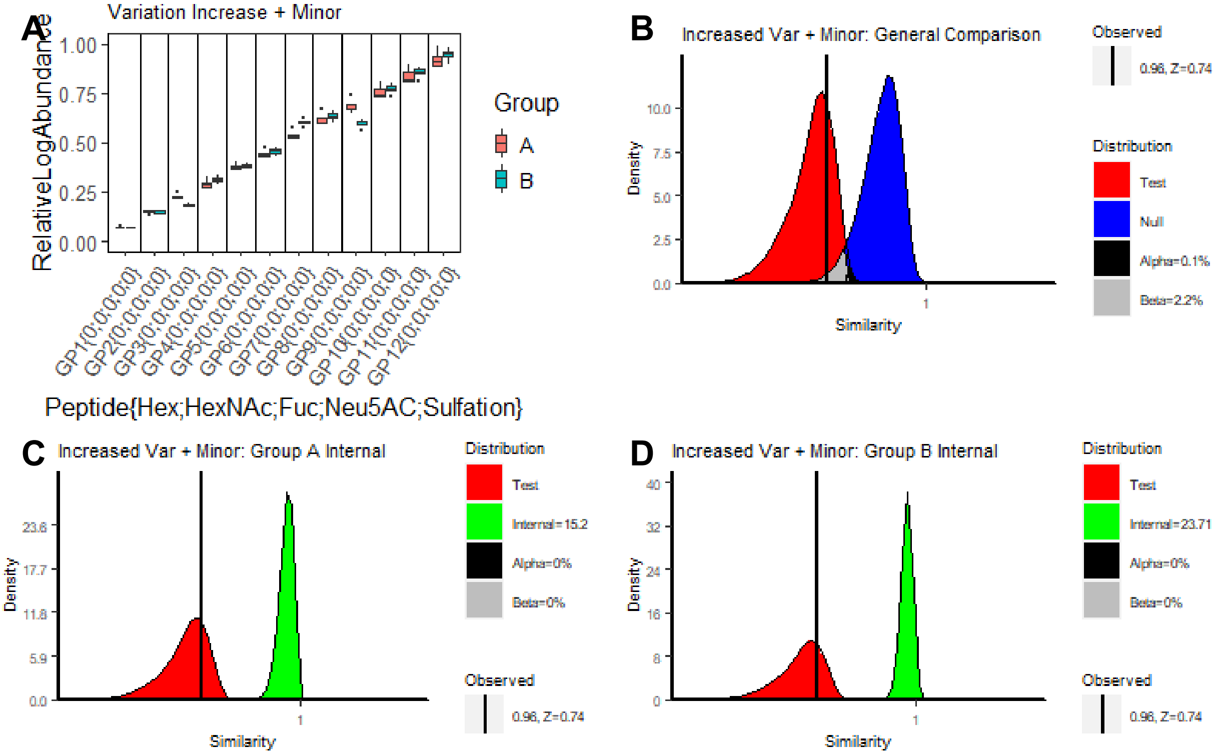

The Internal Similarities, shown in S9c and S9d for completeness, both have ICs (15.2 and 23.71 respectively) greater than 2, and the observed is well simulated by the Test with a z-score of 0.74. The general comparison of Test and Null in S9b shows minimal overlap with a 0.1% FPR and a 2.2% FNR, producing a similar level of confidence in the differentiation between Test and Null as was present in the Major deviation example in Supplemental Figure 8.

In the ranking information, shown in Supplementary Table 4, no identifications exceed significance thresholds as expected. This allows the user to conclude that the overall glycosylation pattern has changed, but individual glycopeptides have too subtle of changes to be reliably discerned by RAMZIS. The highest ranked glycopeptides in this scenario make good candidates for follow up targeted experiments, or for more careful statistical analysis. Regardless of the individual results, the knowledge that there is a detectable disruption to the underlying glycosylation pattern for a glycosite innately informs the user to more closely investigate the discrepancies found.

**Supplemental Table 4: Ranking Minor Deviations**

| Glycopeptides | ZScore | PassOverall | FailCause |
|---|---|---|---|
| GP9 | -1.556410091 | TRUE | Passed |
| GP7 | -1.442600868 | FALSE | LowQualityIn_File1 |
| GP3 | -0.8891118134 | TRUE | Passed |
| GP12 | -0.3878931912 | FALSE | LowQualityIn_File1 |
| GP6 | -0.3522643834 | FALSE | LowQualityIn_File1 |
| GP11 | -0.2977126149 | FALSE | LowQualityIn_File1 |
| GP10 | -0.1721278478 | FALSE | LowQualityIn_File1 |
| GP4 | -0.1426586041 | FALSE | LowQualityIn_File1 |
| GP8 | -0.138142124 | FALSE | LowQualityIn_Both |
| GP5 | -0.09878190344 | FALSE | LowQualityIn_Both |
| GP2 | -0.03038656801 | FALSE | LowQualityIn_Both |
| GP1 | 0.09365809483 | FALSE | LowQualityIn_Both |

#### Theoretical Case: Outlier Identification

RAMZIS identifies sample-wide outliers in sample groups; identification of glycopeptide specific outliers is not a default function of RAMZIS but could theoretically be done using a similar process. RAMZIS should identify samples with: increased rates of missingness, significant shifts in the mean glycosite abundance, mischaracterized samples, and true behavioral outliers. An example of an increased missingness outlier can be found below in the AGP Outlier Removal section at Supplemental Figure []. RAMZIS does not identify what type of outliers it identifies as that requires manual interpretation, but it cannot identify outliers below a specific effect size. It is possible for a sample to be an outlier for one glycosite, but not others, and so the user should not blindly apply outlier removal without further examination. Effect size is directly correlated to RAMZIS' ability to identify outliers, and so subtle outliers may not be identified; individual glycosite outliers can only be identified in extreme cases.

##### Supplemental Figure 10: Theoretical: Outlier Removal

Here in Supplemental Figure 10 we present an example of a global shift in glycosite abundance for an individual sample. The data used is the increased variance data from Supplemental Figure 8 with group B's sample 1 reduced by 10%. Due to the high overall similarity, the default RAMZIS plotting functions were not used to illustrate this example.

S10a and S10d both contain Relative Log Abundance boxplots, S10a is with the 10% shifted sample while S10d is without it. While the effect size is limited, such that the difference is barely discernible in the plots, it is identifiable as shown in S10b and S10c. S10c is output text from RAMZISMain() that informs the user there is likely an outlier. That outlier is identified in S10b, where the text output shows the outlier is the first sample, as it is indexed at [[1]], and that sample 1 is over-represented in the peak at 0.989 with a z-score of 18.7 and under-represented in the peak 0.996 with a z-score of -19.4. These peaks are plotted above this text output in a density plot of the Internal similarity of group B. The dashed lines are the under or over represented peaks in question while the solid lines are the peak minima. After removing the outlier sample in S10d, rerunning RAMZISMain(), we can then make the same kind of plot for S10e with the new comparison. We see that the plot is still multimodal, but the output indicates this comes from sample 4, and that the sample is under-represented in the second highest peak. While it is up to the user's discretion, this outlier was not removed as it was not under-represented in the highest, most-similar peak. This can be thought of as a positive outlier, a sample that is highly consistent. It is worth noting that the emergence of negative outliers, the kind that should be removed, after removal of a different outlier, may indicate the existence of a subgroup rather than a true outlier.

**Supplemental Figure 10: Outlier Removal**

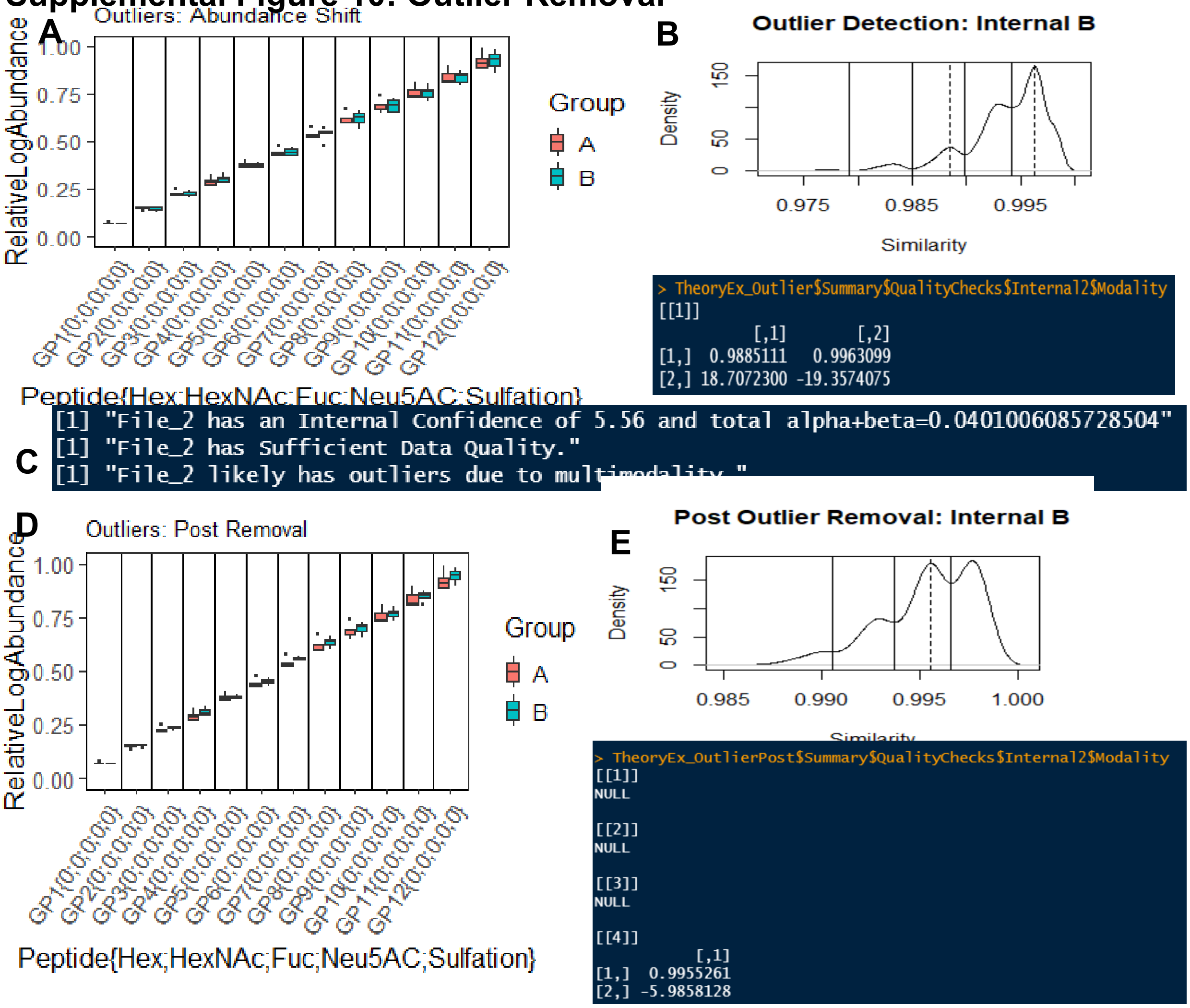

Manual examination of outliers is still critical with this tool. Even in this example, a manual examination of the outlier would reveal its uniform decrease in abundance. Ideally, this should prompt the user to identify the source of this deviation. If it's from a decrease in the protein level associated with that individual sample or if there is an overall decrease in the glycoproteomic signal in that sample, then the answer may not be outlier removal, but signal correction.

### AGP Case-Study: Pure AGP

This section contains further elaboration on the analysis performed in the main text of the article. Outlier Removal refers to the identification and removal of an outlier sample in the Mix 1 AGP data. Full Ranking Summary contains the ranking data of the main-text analysis of Mix 1: Informed vs Naive.

#### Outlier Removal

In Mix 1, Informed vs Naive, there were originally four samples, but one of them was identified as an outlier as shown in Supplemental Figure 11. S11A shows the boxplot of relative log abundances, which shows a much higher degree of missingness compared to the outlier removed dataset in Figure 1A. Figure 1B shows the general comparison of Test and Null similarities, which show an FPR over the 5% threshold at 6%, and an FNR of 17.4%. Visually, the Test and Null similarities have substantially longer left-tails, which would account for much of the increased overlap. RAMZIS still successfully simulated this comparison with an observed similarity at a z-score of 1.25 in the Test similarity.

#### Supplemental Figure 11: AGP: Outlier Removal

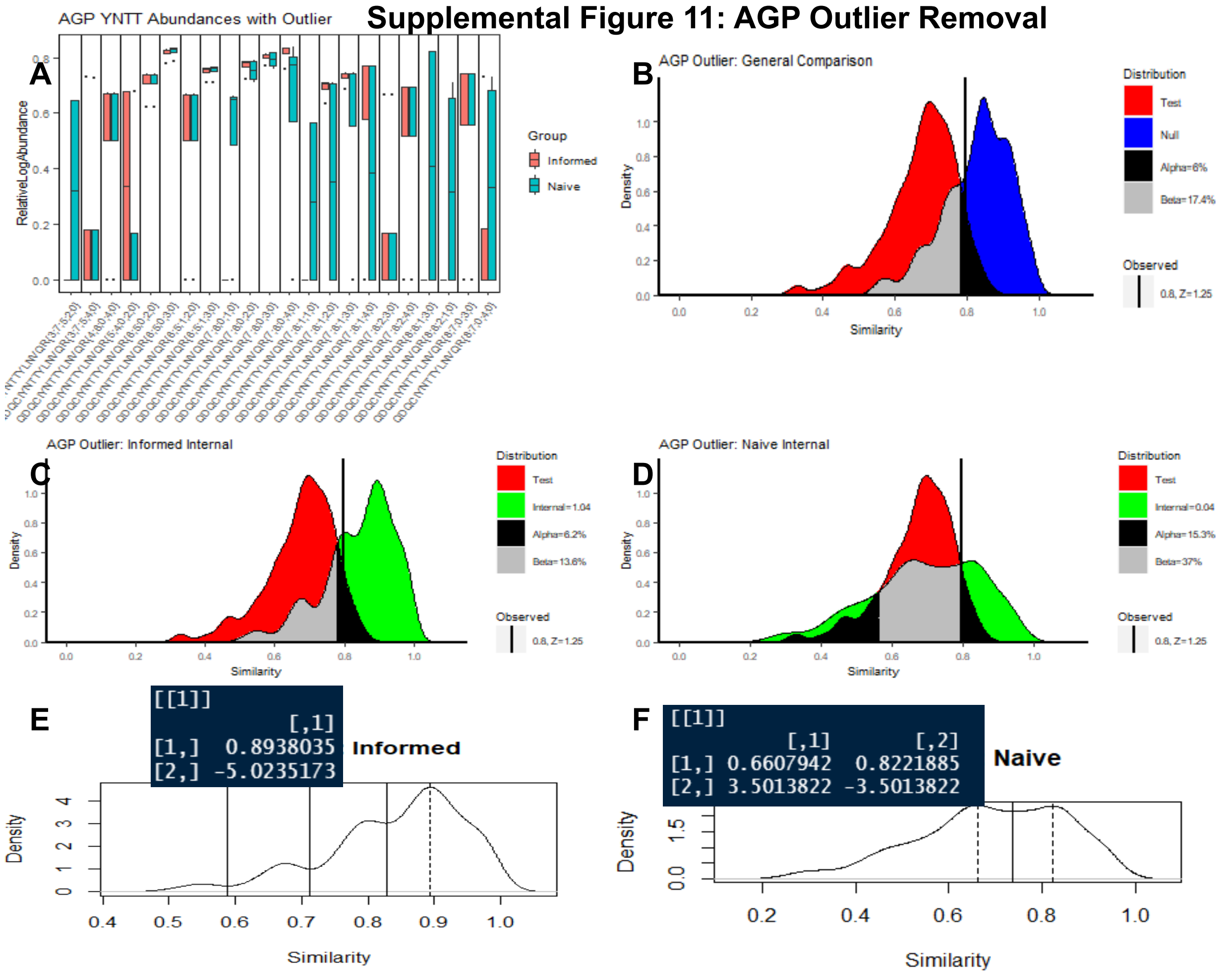

Figure S11C and S11D both show standard RAMZIS Internal similarity outputs, and both Naive and Informed have multimodality ongoing, though the Naive is not substantially different from its outlier removed comparison in Figure 1B. To help illustrate the outliers location, Figures S11E and S11F show the reduced plots found in S10, where the internal similarities are plotted on their own with the pertinent peaks marked with dashed lines and local minima marked with solid lines to show the boundaries of peak ownership. The RAMZIS text output has been overlaid with these plots to show the over and under representation in correspondence with the dashed peaks. In S11E, the Informed search has sample 1 (designated [[1]]) underrepresented (the second row z-score: -5.02) in the peak with a maxima at similarity at 0.894. This peak is the highest, most similar in the Informed Internal similarity, meaning that there are fewer observations of sample 1 in that peak than expected. S11F shows that the Naive search also has an under-representation (z-score=-3.501) in its most similar peak (0.822) and is over-represented (z=3.501) in its secondary peak (0.661).

This information shows that sample 1, in both searches, is an outlier in behavior, and is so in a deleterious way as it is under-represented in the most similar peaks of the Internal similarities.

Further analysis of Sample 1 showed that it had a higher rate of missingness. When it did observe glycopeptides, the abundances were not outside of leave-one-out variances, meaning that its outlier behavior is entirely from its missingness. The TIC from this run was substantially lower than the other run's TICs, indicating that there was likely a sample preparation or LC related error of some kind resulting in lower signal strength.

#### Full Ranking Summary

This section contains the full glycopeptide ranking table from the analysis performed in section 3.1.1: Naive vs Informed search spaces in pure AGP after outlier removal. This can be found in Supplemental Table 5.

##### Supplemental Table 5: Pure AGP Glycopeptide Rankings

| Glycopeptides | ZScore | PassOverall | FailCause |
|---|---|---|---|
| QDQCIYNTTYLNVQR{Hex:7; HexNAc:6; Neu5Ac:1} | -1.946552562 | FALSE | UnseenIn_File2 |
| QDQCIYNTTYLNVQR{Hex:6; HexNAc:5; Neu5Ac:3} | -0.9344698552 | TRUE | Passed |
| QDQCIYNTTYLNVQR{Fuc:1; Hex:7; HexNAc:6; Neu5Ac:1} | -0.8863841745 | FALSE | LowQualityIn_Both |
| QDQCIYNTTYLNVQR{@sulfate:1; Fuc:5; Hex:3; HexNAc:7; Neu5Ac:2} | -0.8849453208 | FALSE | LowQualityIn_Both |
| QDQCIYNTTYLNVQR{Fuc:1; Hex:8; HexNAc:6; Neu5Ac:3} | -0.880557342 | FALSE | LowQualityIn_Both |
| QDQCIYNTTYLNVQR{Fuc:1; Hex:6; HexNAc:5; Neu5Ac:3} | -0.6147228118 | TRUE | Passed |
| QDQCIYNTTYLNVQR{Hex:7; HexNAc:6; Neu5Ac:4} | -0.4792286414 | TRUE | Passed |
| QDQCIYNTTYLNVQR{Hex:7; HexNAc:6; Neu5Ac:2} | -0.1444044722 | TRUE | Passed |
| QDQCIYNTTYLNVQR{Hex:5; HexNAc:4; Neu5Ac:2} | -0.1323206534 | FALSE | LowQualityIn_Both |
| QDQCIYNTTYLNVQR{Hex:8; HexNAc:7; Neu5Ac:4} | -0.1174572186 | FALSE | LowQualityIn_Both |
| QDQCIYNTTYLNVQR{Hex:7; HexNAc:6; Neu5Ac:3} | -0.1106629259 | TRUE | Passed |
| QDQCIYNTTYLNVQR{Fuc:1; Hex:7; HexNAc:6; Neu5Ac:2} | -0.09310602893 | FALSE | LowQualityIn_File1 |
| QDQCIYNTTYLNVQR{Fuc:1; Hex:7; HexNAc:6; Neu5Ac:4} | -0.09155135654 | FALSE | LowQualityIn_File1 |
| QDQCIYNTTYLNVQR{Fuc:1; Hex:7; HexNAc:6; Neu5Ac:3} | -0.05578959387 | TRUE | Passed |
| QDQCIYNTTYLNVQR{Hex:6; HexNAc:5; Neu5Ac:2} | -0.05196458121 | TRUE | Passed |
| QDQCIYNTTYLNVQR{Hex:8; HexNAc:7; Neu5Ac:3} | -0.04765218453 | TRUE | Passed |
| QDQCIYNTTYLNVQR{@sulfate:1; Hex:4; HexNAc:6; Neu5Ac:4} | -0.04729701855 | TRUE | Passed |
| QDQCIYNTTYLNVQR{Fuc:2; Hex:7; HexNAc:6; Neu5Ac:4} | -0.04723632322 | TRUE | Passed |
| QDQCIYNTTYLNVQR{Fuc:1; Hex:6; HexNAc:5; Neu5Ac:2} | -0.04632468883 | TRUE | Passed |
| QDQCIYNTTYLNVQR{Fuc:2; Hex:7; HexNAc:6; Neu5Ac:3} | -0.007826979993 | FALSE | LowQualityIn_Both |
| QDQCIYNTTYLNVQR{@sulfate:1; Fuc:5; Hex:3; HexNAc:7; Neu5Ac:4} | -0.001148441625 | FALSE | LowQualityIn_Both |
| QDQCIYNTTYLNVQR{@sulfate:1; Fuc:2; Hex:4; HexNAc:8; Neu5Ac:1} | NA | FALSE | BelowMinimumObs |
| QDQCIYNTTYLNVQR{@sulfate:1; Fuc:2; Hex:4; HexNAc:8; Neu5Ac:3} | NA | FALSE | BelowMinimumObs |
| QDQCIYNTTYLNVQR{@sulfate:1; Hex:4; HexNAc:6; Neu5Ac:3} | NA | FALSE | BelowMinimumObs |
| QDQCIYNTTYLNVQR{Fuc:1; Hex:6; HexNAc:4; Neu5Ac:1} | NA | FALSE | BelowMinimumObs |
| QDQCIYNTTYLNVQR{Fuc:1; Hex:8; HexNAc:6; Neu5Ac:1} | NA | FALSE | BelowMinimumObs |
| QDQCIYNTTYLNVQR{Fuc:1; Hex:8; HexNAc:6; Neu5Ac:2} | NA | FALSE | BelowMinimumObs |
| QDQCIYNTTYLNVQR{Fuc:1; Hex:8; HexNAc:7; Neu5Ac:4} | NA | FALSE | BelowMinimumObs |
| QDQCIYNTTYLNVQR{Fuc:2; Hex:8; HexNAc:6; Neu5Ac:1} | NA | FALSE | BelowMinimumObs |
| QDQCIYNTTYLNVQR{Fuc:2; Hex:8; HexNAc:6; Neu5Ac:3} | NA | FALSE | BelowMinimumObs |
| QDQCIYNTTYLNVQR{Hex:8; HexNAc:7; Neu5Ac:2} | NA | FALSE | BelowMinimumObs |

### AGP Case-Study: Mix 1 vs Mix 5

This section provides additional context to the analysis pertaining to section 3.1.2 and Figure 2, the comparisons of Mix 1 (pure AGP) and Mix 5 in both the Naive and Informed comparisons with respect to glycosite YnTT.

#### Naive Scaled

The rankings in Supplemental Table 6 pertain to the scaled analysis done in 3.1.2. Scaling was done to account for a global shift in abundance observed in Mix 5. Naive Mix 5 was scaled by 1.067.

##### Full Ranking Summary

##### Supplemental Table 6: Naive: Mix 1 vs Mix 5 Scaled

| Glycopeptides | ZScore | PassOverall | FailCause |
|---|---|---|---|
| QDQCIYNTTYLNVQR{Hex:7; HexNAc:6; Neu5Ac:1} | -2.017354535 | FALSE | UnseenIn_File2 |
| QDQCIYNTTYLNVQR{Fuc:1; Hex:6; HexNAc:5; Neu5Ac:2} | -2.01515825 | FALSE | UnseenIn_File2 |
| QDQCIYNTTYLNVQR{@sulfate:1; Hex:4; HexNAc:6; Neu5Ac:4} | -2.014661812 | FALSE | UnseenIn_File2 |
| QDQCIYNTTYLNVQR{Fuc:2; Hex:7; HexNAc:6; Neu5Ac:4} | -2.00870877 | FALSE | UnseenIn_File2 |
| QDQCIYNTTYLNVQR{Fuc:1; Hex:7; HexNAc:6; Neu5Ac:3} | -2.000440825 | FALSE | UnseenIn_File2 |
| QDQCIYNTTYLNVQR{Hex:8; HexNAc:7; Neu5Ac:3} | -1.998416057 | FALSE | UnseenIn_File2 |
| QDQCIYNTTYLNVQR{Fuc:1; Hex:6; HexNAc:5; Neu5Ac:3} | -1.994785183 | FALSE | UnseenIn_File2 |
| QDQCIYNTTYLNVQR{@sulfate:1; Hex:4; HexNAc:8; Neu5Ac:3} | -1.993957069 | FALSE | UnseenIn_File1 |
| QDQCIYNTTYLNVQR{Hex:6; HexNAc:5; Neu5Ac:3} | -1.000902286 | FALSE | LowQualityIn_File1 |
| QDQCIYNTTYLNVQR{Hex:6; HexNAc:5; Neu5Ac:2} | -0.9274038643 | FALSE | LowQualityIn_File1 |
| QDQCIYNTTYLNVQR{Fuc:1; Hex:7; HexNAc:6; Neu5Ac:1} | -0.8863143119 | FALSE | LowQualityIn_Both |
| QDQCIYNTTYLNVQR{@sulfate:1; Fuc:5; Hex:3; HexNAc:7; Neu5Ac:2} | -0.8849555986 | FALSE | LowQualityIn_Both |
| QDQCIYNTTYLNVQR{@sulfate:1; Fuc:5; Hex:3; HexNAc:7; Neu5Ac:3} | -0.8794608999 | FALSE | LowQualityIn_Both |
| QDQCIYNTTYLNVQR{@sulfate:1; Fuc:1; Hex:4; HexNAc:8; Neu5Ac:3} | -0.8792374146 | FALSE | LowQualityIn_Both |
| QDQCIYNTTYLNVQR{@sulfate:1; Fuc:1; Hex:5; HexNAc:8; Neu5Ac:2} | -0.8790271722 | FALSE | LowQualityIn_Both |
| QDQCIYNTTYLNVQR{Fuc:1; Hex:7; HexNAc:6; Neu5Ac:2} | -0.8566172218 | FALSE | LowQualityIn_Both |
| QDQCIYNTTYLNVQR{Hex:7; HexNAc:6; Neu5Ac:3} | -0.5715212188 | FALSE | LowQualityIn_File2 |
| QDQCIYNTTYLNVQR{Fuc:1; Hex:8; HexNAc:6; Neu5Ac:3} | -0.1200502352 | FALSE | LowQualityIn_Both |
| QDQCIYNTTYLNVQR{Hex:7; HexNAc:6; Neu5Ac:2} | -0.1054248179 | FALSE | LowQualityIn_File1 |
| QDQCIYNTTYLNVQR{Hex:7; HexNAc:6; Neu5Ac:4} | -0.07346798053 | FALSE | LowQualityIn_File2 |
| QDQCIYNTTYLNVQR{Fuc:2; Hex:8; HexNAc:6; Neu5Ac:3} | -0.0112437359 | FALSE | LowQualityIn_Both |
| QDQCIYNTTYLNVQR{Fuc:1; Hex:6; HexNAc:4; Neu5Ac:1} | 0.0006557152411 | FALSE | LowQualityIn_Both |
| QDQCIYNTTYLNVQR{Fuc:1; Hex:7; HexNAc:6; Neu5Ac:4} | 0.007203044013 | FALSE | LowQualityIn_Both |
| QDQCIYNTTYLNVQR{Hex:8; HexNAc:7; Neu5Ac:4} | 0.009189220675 | FALSE | LowQualityIn_Both |
| QDQCIYNTTYLNVQR{@sulfate:1; Fuc:1; Hex:3; HexNAc:7; Neu5Ac:3} | NA | FALSE | BelowMinimumObs |
| QDQCIYNTTYLNVQR{@sulfate:1; Fuc:2; Hex:4; HexNAc:7; Neu5Ac:2} | NA | FALSE | BelowMinimumObs |
| QDQCIYNTTYLNVQR{@sulfate:1; Fuc:2; Hex:4; HexNAc:8; Neu5Ac:1} | NA | FALSE | BelowMinimumObs |
| QDQCIYNTTYLNVQR{@sulfate:1; Fuc:2; Hex:5; HexNAc:8; Neu5Ac:2} | NA | FALSE | BelowMinimumObs |
| QDQCIYNTTYLNVQR{@sulfate:1; Fuc:5; Hex:3; HexNAc:7; Neu5Ac:4} | NA | FALSE | BelowMinimumObs |
| QDQCIYNTTYLNVQR{@sulfate:1; Hex:10; HexNAc:5; Neu5Ac:1} | NA | FALSE | BelowMinimumObs |
| QDQCIYNTTYLNVQR{@sulfate:1; Hex:3; HexNAc:7; Neu5Ac:3} | NA | FALSE | BelowMinimumObs |
| QDQCIYNTTYLNVQR{@sulfate:1; Hex:4; HexNAc:6; Neu5Ac:3} | NA | FALSE | BelowMinimumObs |
| QDQCIYNTTYLNVQR{Fuc:1; Hex:8; HexNAc:6; Neu5Ac:1} | NA | FALSE | BelowMinimumObs |
| QDQCIYNTTYLNVQR{Fuc:1; Hex:8; HexNAc:6; Neu5Ac:2} | NA | FALSE | BelowMinimumObs |
| QDQCIYNTTYLNVQR{Fuc:1; Hex:8; HexNAc:7; Neu5Ac:4} | NA | FALSE | BelowMinimumObs |
| QDQCIYNTTYLNVQR{Fuc:2; Hex:7; HexNAc:5; Neu5Ac:2} | NA | FALSE | BelowMinimumObs |
| QDQCIYNTTYLNVQR{Fuc:2; Hex:7; HexNAc:6; Neu5Ac:3} | NA | FALSE | BelowMinimumObs |
| QDQCIYNTTYLNVQR{Fuc:2; Hex:8; HexNAc:6; Neu5Ac:1} | NA | FALSE | BelowMinimumObs |
| QDQCIYNTTYLNVQR{Fuc:2; Hex:8; HexNAc:6; Neu5Ac:2} | NA | FALSE | BelowMinimumObs |
| QDQCIYNTTYLNVQR{Hex:5; HexNAc:4; Neu5Ac:2} | NA | FALSE | BelowMinimumObs |
| QDQCIYNTTYLNVQR{Hex:8; HexNAc:7; Neu5Ac:2} | NA | FALSE | BelowMinimumObs |

#### Naive Unscaled

##### Similarity Analysis

For completeness, the unscaled analysis is provided here in brief in Supplemental Figure 12. S12A contains the abundance boxplot, where there is a pronounced scalar shift. S12B and S12C show the Internal similarities of Mix 1 and 5 respectively which have ICs of 4.1 and 1.63 respectively. The observed similarity is well-simulated with a z-score of 0.87, and there is a clear separation between the Test and Null similarities as seen in S12D, the general comparison of Test and Null. S12E shows the linear model used to determine the scaling value used in the main text analysis. None of this analysis substantially differs from the Naive analysis of the main text.

##### Supplemental Figure 12: AGP: Naive Unscaled: Mix 1 vs Mix 5

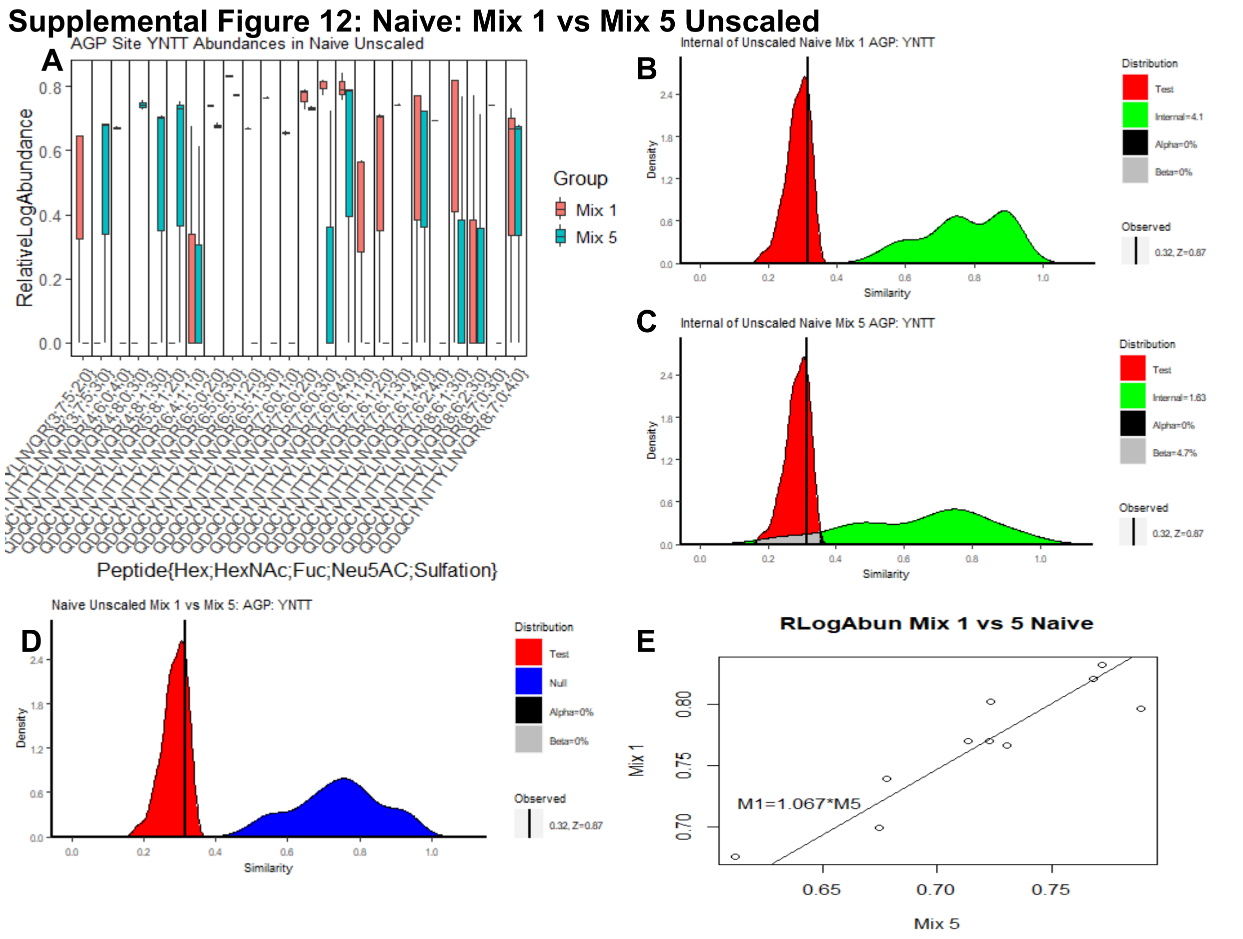

##### Full Ranking Summary

For completeness, the rankings of the unscaled Naive analysis are provided here. The order of glycopeptides is shifted, and several z-scores have dropped below significance, but the primary cause of differentiation in the highest ranked contributors to dissimilarity is the same as the scaled: disjoint observations between the two sample groups.

**Supplemental Table 7: Naive: Mix 1 vs Mix 5 Unscaled**

| Glycopeptides | ZScore | PassOverall | FailCause |
|---|---|---|---|
| QDQCIYNTTYLNVQR{Hex:7; HexNAc:6; Neu5Ac:1} | -1.978072065 | FALSE | UnseenIn_File2 |
| QDQCIYNTTYLNVQR{Fuc:1; Hex:6; HexNAc:5; Neu5Ac:2} | -1.974507167 | FALSE | UnseenIn_File2 |
| QDQCIYNTTYLNVQR{@sulfate:1; Hex:4; HexNAc:6; Neu5Ac:4} | -1.973847799 | FALSE | UnseenIn_File2 |
| QDQCIYNTTYLNVQR{Fuc:2; Hex:7; HexNAc:6; Neu5Ac:4} | -1.966647442 | FALSE | UnseenIn_File2 |
| QDQCIYNTTYLNVQR{@sulfate:1; Hex:4; HexNAc:8; Neu5Ac:3} | -1.960116619 | FALSE | UnseenIn_File1 |
| QDQCIYNTTYLNVQR{Fuc:1; Hex:7; HexNAc:6; Neu5Ac:3} | -1.958942465 | FALSE | UnseenIn_File2 |
| QDQCIYNTTYLNVQR{Hex:8; HexNAc:7; Neu5Ac:3} | -1.957859691 | FALSE | UnseenIn_File2 |
| QDQCIYNTTYLNVQR{Fuc:1; Hex:6; HexNAc:5; Neu5Ac:3} | -1.95526794 | FALSE | UnseenIn_File2 |
| QDQCIYNTTYLNVQR{Hex:6; HexNAc:5; Neu5Ac:3} | -1.831499753 | FALSE | LowQualityIn_File1 |
| QDQCIYNTTYLNVQR{Hex:6; HexNAc:5; Neu5Ac:2} | -1.596834036 | FALSE | LowQualityIn_File1 |
| QDQCIYNTTYLNVQR{@sulfate:1; Fuc:1; Hex:5; HexNAc:8; Neu5Ac:2} | -0.8931595811 | FALSE | LowQualityIn_Both |
| QDQCIYNTTYLNVQR{@sulfate:1; Fuc:1; Hex:4; HexNAc:8; Neu5Ac:3} | -0.8926321709 | FALSE | LowQualityIn_Both |
| QDQCIYNTTYLNVQR{@sulfate:1; Fuc:5; Hex:3; HexNAc:7; Neu5Ac:3} | -0.8923796016 | FALSE | LowQualityIn_Both |
| QDQCIYNTTYLNVQR{Fuc:1; Hex:7; HexNAc:6; Neu5Ac:2} | -0.8826370706 | FALSE | LowQualityIn_Both |
| QDQCIYNTTYLNVQR{Fuc:1; Hex:7; HexNAc:6; Neu5Ac:1} | -0.8348766778 | FALSE | LowQualityIn_Both |
| QDQCIYNTTYLNVQR{@sulfate:1; Fuc:5; Hex:3; HexNAc:7; Neu5Ac:2} | -0.8333030862 | FALSE | LowQualityIn_Both |
| QDQCIYNTTYLNVQR{Hex:7; HexNAc:6; Neu5Ac:3} | -0.6642829836 | FALSE | LowQualityIn_File2 |
| QDQCIYNTTYLNVQR{Hex:7; HexNAc:6; Neu5Ac:2} | -0.545866928 | FALSE | LowQualityIn_File1 |
| QDQCIYNTTYLNVQR{Fuc:1; Hex:8; HexNAc:6; Neu5Ac:3} | -0.1719883895 | FALSE | LowQualityIn_Both |
| QDQCIYNTTYLNVQR{Hex:7; HexNAc:6; Neu5Ac:4} | -0.1044163703 | FALSE | LowQualityIn_File2 |
| QDQCIYNTTYLNVQR{Fuc:1; Hex:6; HexNAc:4; Neu5Ac:1} | -0.01235929583 | FALSE | LowQualityIn_Both |
| QDQCIYNTTYLNVQR{Fuc:2; Hex:8; HexNAc:6; Neu5Ac:3} | -0.01192585312 | FALSE | LowQualityIn_Both |
| QDQCIYNTTYLNVQR{Fuc:1; Hex:7; HexNAc:6; Neu5Ac:4} | -0.008072765172 | FALSE | LowQualityIn_Both |
| QDQCIYNTTYLNVQR{Hex:8; HexNAc:7; Neu5Ac:4} | 0.001870856783 | FALSE | LowQualityIn_Both |
| QDQCIYNTTYLNVQR{@sulfate:1; Fuc:1; Hex:3; HexNAc:7; Neu5Ac:3} | NA | FALSE | BelowMinimumObs |
| QDQCIYNTTYLNVQR{@sulfate:1; Fuc:2; Hex:4; HexNAc:7; Neu5Ac:2} | NA | FALSE | BelowMinimumObs |
| QDQCIYNTTYLNVQR{@sulfate:1; Fuc:2; Hex:4; HexNAc:8; Neu5Ac:1} | NA | FALSE | BelowMinimumObs |
| QDQCIYNTTYLNVQR{@sulfate:1; Fuc:2; Hex:5; HexNAc:8; Neu5Ac:2} | NA | FALSE | BelowMinimumObs |
| QDQCIYNTTYLNVQR{@sulfate:1; Fuc:5; Hex:3; HexNAc:7; Neu5Ac:4} | NA | FALSE | BelowMinimumObs |
| QDQCIYNTTYLNVQR{@sulfate:1; Hex:10; HexNAc:5; Neu5Ac:1} | NA | FALSE | BelowMinimumObs |
| QDQCIYNTTYLNVQR{@sulfate:1; Hex:3; HexNAc:7; Neu5Ac:3} | NA | FALSE | BelowMinimumObs |
| QDQCIYNTTYLNVQR{@sulfate:1; Hex:4; HexNAc:6; Neu5Ac:3} | NA | FALSE | BelowMinimumObs |
| QDQCIYNTTYLNVQR{Fuc:1; Hex:8; HexNAc:6; Neu5Ac:1} | NA | FALSE | BelowMinimumObs |
| QDQCIYNTTYLNVQR{Fuc:1; Hex:8; HexNAc:6; Neu5Ac:2} | NA | FALSE | BelowMinimumObs |
| QDQCIYNTTYLNVQR{Fuc:1; Hex:8; HexNAc:7; Neu5Ac:4} | NA | FALSE | BelowMinimumObs |
| QDQCIYNTTYLNVQR{Fuc:2; Hex:7; HexNAc:5; Neu5Ac:2} | NA | FALSE | BelowMinimumObs |
| QDQCIYNTTYLNVQR{Fuc:2; Hex:7; HexNAc:6; Neu5Ac:3} | NA | FALSE | BelowMinimumObs |
| QDQCIYNTTYLNVQR{Fuc:2; Hex:8; HexNAc:6; Neu5Ac:1} | NA | FALSE | BelowMinimumObs |
| QDQCIYNTTYLNVQR{Fuc:2; Hex:8; HexNAc:6; Neu5Ac:2} | NA | FALSE | BelowMinimumObs |
| QDQCIYNTTYLNVQR{Hex:5; HexNAc:4; Neu5Ac:2} | NA | FALSE | BelowMinimumObs |
| QDQCIYNTTYLNVQR{Hex:8; HexNAc:7; Neu5Ac:2} | NA | FALSE | BelowMinimumObs |

#### Informed

The rankings in Supplemental Table 8 pertain to the scaled analysis done in 3.1.2. Scaling was done to account for a global shift in abundance observed in Mix 5. Informed Mix 5 was scaled by 1.084.

#### Full Ranking Summary

##### Supplemental Table 8: Informed: Mix 1 vs Mix 5 Scaled

| Glycopeptides | ZScore | PassOverall | FailCause |
|---|---|---|---|
| QDQCIYNTTYLNVQR{Fuc:1; Hex:6; HexNAc:5; Neu5Ac:2} | -2.016914538 | FALSE | UnseenIn_File2 |
| QDQCIYNTTYLNVQR{@sulfate:1; Hex:4; HexNAc:6; Neu5Ac:4} | -2.016374361 | FALSE | UnseenIn_File2 |
| QDQCIYNTTYLNVQR{Fuc:1; Hex:7; HexNAc:6; Neu5Ac:2} | -2.011076058 | FALSE | UnseenIn_File2 |
| QDQCIYNTTYLNVQR{Fuc:2; Hex:7; HexNAc:6; Neu5Ac:4} | -2.007856868 | FALSE | UnseenIn_File2 |
| QDQCIYNTTYLNVQR{Fuc:1; Hex:7; HexNAc:6; Neu5Ac:3} | -2.000628783 | FALSE | UnseenIn_File2 |
| QDQCIYNTTYLNVQR{Hex:8; HexNAc:7; Neu5Ac:3} | -1.998418323 | FALSE | UnseenIn_File2 |
| QDQCIYNTTYLNVQR{@sulfate:1; Fuc:1; Hex:3; HexNAc:7; Neu5Ac:3} | -1.996835746 | FALSE | UnseenIn_File1 |
| QDQCIYNTTYLNVQR{@sulfate:1; Fuc:1; Hex:4; HexNAc:8; Neu5Ac:3} | -1.9952286 | FALSE | UnseenIn_File1 |
| QDQCIYNTTYLNVQR{@sulfate:1; Hex:4; HexNAc:8; Neu5Ac:3} | -1.987394794 | FALSE | UnseenIn_File1 |
| QDQCIYNTTYLNVQR{Hex:6; HexNAc:5; Neu5Ac:3} | -1.132543698 | TRUE | Passed |
| QDQCIYNTTYLNVQR{Fuc:1; Hex:6; HexNAc:5; Neu5Ac:3} | -0.8417743956 | FALSE | LowQualityIn_File2 |
| QDQCIYNTTYLNVQR{Hex:7; HexNAc:6; Neu5Ac:2} | -0.6213296245 | TRUE | Passed |
| QDQCIYNTTYLNVQR{Hex:7; HexNAc:6; Neu5Ac:3} | -0.5665884601 | FALSE | LowQualityIn_File2 |
| QDQCIYNTTYLNVQR{Fuc:1; Hex:7; HexNAc:6; Neu5Ac:4} | -0.4973306375 | FALSE | LowQualityIn_File2 |
| QDQCIYNTTYLNVQR{Hex:7; HexNAc:6; Neu5Ac:4} | -0.3426218304 | TRUE | Passed |
| QDQCIYNTTYLNVQR{Hex:6; HexNAc:5; Neu5Ac:2} | -0.2106545612 | TRUE | Passed |
| QDQCIYNTTYLNVQR{Hex:5; HexNAc:4; Neu5Ac:2} | -0.1396757738 | FALSE | LowQualityIn_Both |
| QDQCIYNTTYLNVQR{@sulfate:1; Fuc:2; Hex:4; HexNAc:8; Neu5Ac:3} | NA | FALSE | BelowMinimumObs |
| QDQCIYNTTYLNVQR{@sulfate:1; Fuc:5; Hex:3; HexNAc:7; Neu5Ac:3} | NA | FALSE | BelowMinimumObs |
| QDQCIYNTTYLNVQR{@sulfate:1; Fuc:5; Hex:3; HexNAc:7; Neu5Ac:4} | NA | FALSE | BelowMinimumObs |
| QDQCIYNTTYLNVQR{@sulfate:1; Hex:3; HexNAc:7; Neu5Ac:3} | NA | FALSE | BelowMinimumObs |
| QDQCIYNTTYLNVQR{Fuc:2; Hex:7; HexNAc:6; Neu5Ac:3} | NA | FALSE | BelowMinimumObs |
| QDQCIYNTTYLNVQR{Hex:8; HexNAc:7; Neu5Ac:4} | NA | FALSE | BelowMinimumObs |

### Informed Unscaled

#### Similarity Analysis

For completeness, the unscaled Informed analysis is provided in Supplemental Figure 13. S13A contains the abundance boxplot, where there is a pronounced scalar shift, more apparent here than in S12A. S13B and S13C show the Internal similarities of Mix 1 and 5 respectively which have ICs of 12.63 and 5.81 respectively. The observed similarity is well-simulated with a z-score of 0.1, and there is a clear separation between the Test and Null similarities as seen in S13D, the general comparison of Test and Null for unscaled Informed. S13E shows the linear model used to determine the scaling value used in the main text analysis. None of this analysis substantially differs from the Informed analysis of the main text.

#### Supplemental Figure 13: AGP: Informed Unscaled: Mix 1 vs Mix 5

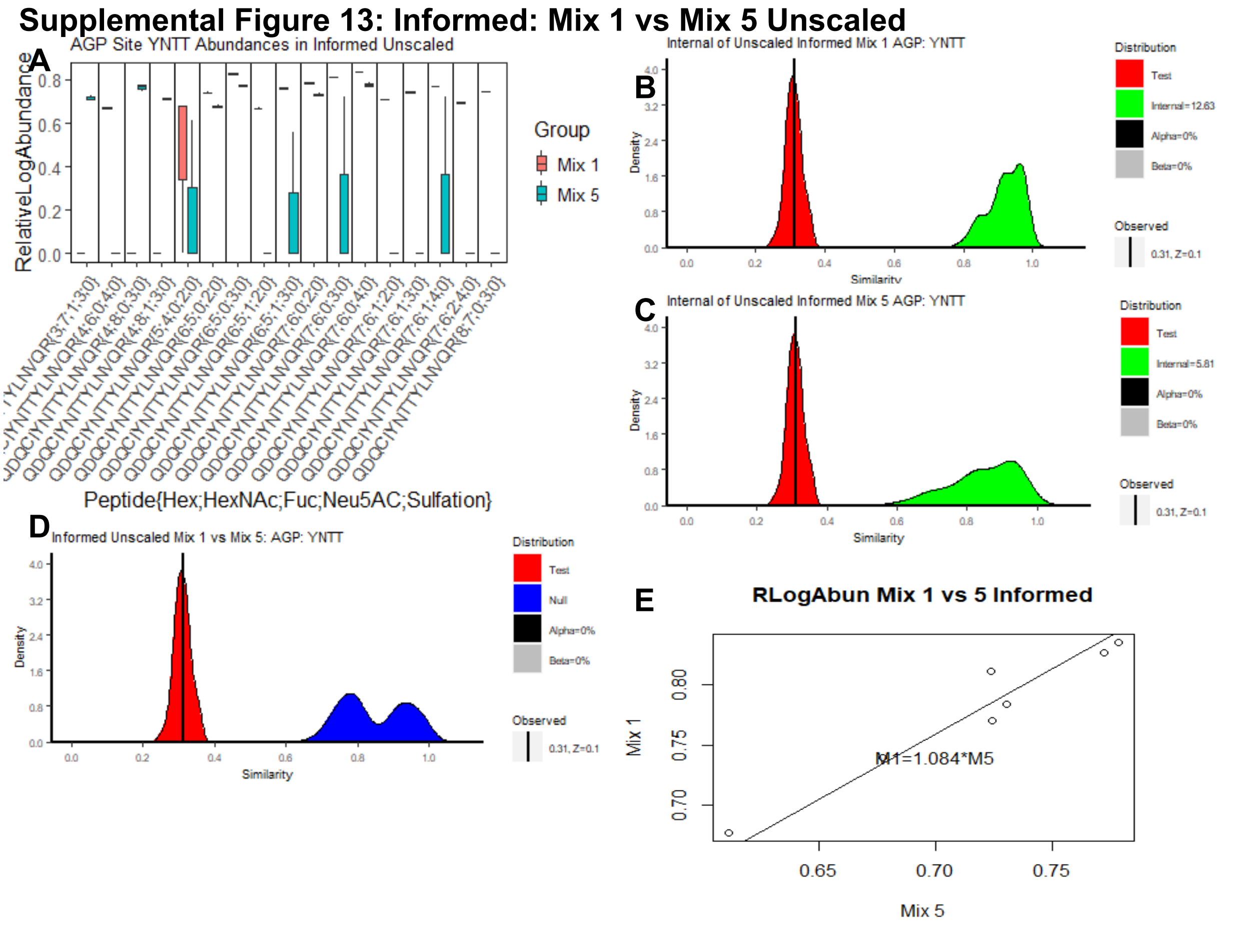

#### Full Ranking Summary

For completeness, the rankings of the unscaled Informed analysis are provided here. The order of glycopeptides is shifted, and several z-scores have dropped below significance, but the primary cause of differentiation in the highest ranked contributors to dissimilarity is once again the same as the scaled analysis: disjoint observations between the two sample groups.

#### Supplemental Table 9: Informed: Mix 1 vs Mix 5 Unscaled

| Glycopeptides | ZScore | PassOverall | FailCause |
|---|---|---|---|
| QDQCIYNTTYLNVQR{Fuc:1; Hex:6; HexNAc:5; Neu5Ac:2} | -1.949874116 | FALSE | UnseenIn_File2 |
| QDQCIYNTTYLNVQR{@sulfate:1; Hex:4; HexNAc:6; Neu5Ac:4} | -1.94942564 | FALSE | UnseenIn_File2 |
| QDQCIYNTTYLNVQR{@sulfate:1; Fuc:1; Hex:3; HexNAc:7; Neu5Ac:3} | -1.945930146 | FALSE | UnseenIn_File1 |
| QDQCIYNTTYLNVQR{Fuc:2; Hex:7; HexNAc:6; Neu5Ac:4} | -1.944599675 | FALSE | UnseenIn_File2 |
| QDQCIYNTTYLNVQR{@sulfate:1; Fuc:1; Hex:4; HexNAc:8; Neu5Ac:3} | -1.943812194 | FALSE | UnseenIn_File1 |
| QDQCIYNTTYLNVQR{Fuc:1; Hex:7; HexNAc:6; Neu5Ac:2} | -1.943716071 | FALSE | UnseenIn_File2 |
| QDQCIYNTTYLNVQR{@sulfate:1; Hex:4; HexNAc:8; Neu5Ac:3} | -1.939871946 | FALSE | UnseenIn_File1 |
| QDQCIYNTTYLNVQR{Fuc:1; Hex:7; HexNAc:6; Neu5Ac:3} | -1.937281158 | FALSE | UnseenIn_File2 |
| QDQCIYNTTYLNVQR{Hex:8; HexNAc:7; Neu5Ac:3} | -1.935589928 | FALSE | UnseenIn_File2 |
| QDQCIYNTTYLNVQR{Hex:6; HexNAc:5; Neu5Ac:3} | -1.784694778 | TRUE | Passed |
| QDQCIYNTTYLNVQR{Hex:7; HexNAc:6; Neu5Ac:2} | -1.661444459 | TRUE | Passed |

| | | | |
|---|---|---|---|
| QDQCIYNTTYLNVQR{Hex:7; HexNAc:6; Neu5Ac:4} | -1.613022448 | TRUE | Passed |
| QDQCIYNTTYLNVQR{Hex:6; HexNAc:5; Neu5Ac:2} | -1.565836679 | TRUE | Passed |
| QDQCIYNTTYLNVQR{Fuc:1; Hex:6; HexNAc:5; Neu5Ac:3} | -0.9507001545 | FALSE | LowQualityIn_File2 |
| QDQCIYNTTYLNVQR{Hex:7; HexNAc:6; Neu5Ac:3} | -0.6830903983 | FALSE | LowQualityIn_File2 |
| QDQCIYNTTYLNVQR{Fuc:1; Hex:7; HexNAc:6; Neu5Ac:4} | -0.6117663593 | FALSE | LowQualityIn_File2 |
| QDQCIYNTTYLNVQR{Hex:5; HexNAc:4; Neu5Ac:2} | -0.1850272419 | FALSE | LowQualityIn_Both |
| QDQCIYNTTYLNVQR{@sulfate:1; Fuc:2; Hex:4; HexNAc:8; Neu5Ac:3} | NA | FALSE | BelowMinimumObs |
| QDQCIYNTTYLNVQR{@sulfate:1; Fuc:5; Hex:3; HexNAc:7; Neu5Ac:3} | NA | FALSE | BelowMinimumObs |
| QDQCIYNTTYLNVQR{@sulfate:1; Fuc:5; Hex:3; HexNAc:7; Neu5Ac:4} | NA | FALSE | BelowMinimumObs |
| QDQCIYNTTYLNVQR{@sulfate:1; Hex:3; HexNAc:7; Neu5Ac:3} | NA | FALSE | BelowMinimumObs |
| QDQCIYNTTYLNVQR{Fuc:2; Hex:7; HexNAc:6; Neu5Ac:3} | NA | FALSE | BelowMinimumObs |
| QDQCIYNTTYLNVQR{Hex:8; HexNAc:7; Neu5Ac:4} | NA | FALSE | BelowMinimumObs |
